# *Nasonia vitripennis* males exhibit greater effort and competency in detecting hosts with conspecific females than other *Nasonia* males

**DOI:** 10.1101/2025.09.17.676707

**Authors:** Taruna Verma, Bharat Kumar Sirasva, Satyajit Jena, Diptimayee Behera, Ambili Anoop, Ruchira Sen, Rhitoban Raychoudhury

## Abstract

*Nasonia* is a species complex of four parasitoid wasps. *N. vitripennis* is cosmopolitan, while the other three species are micro-sympatric with it. This distribution can select for distinct species-specific mate recognition capabilities. However, whether *Nasonia* males can identify hosts with conspecific females against hosts with heterospecific females is not known. Therefore, we tested this hypothesis in a cafeteria-based choice assay and show that *N. vitripennis* males can distinguish hosts with conspecific wasps against those parasitised by *N. giraulti* and *N. oneida,* exhibiting longer search time and distance traversed with faster search speed. We also found that *N. longicornis* males can distinguish hosts with conspecific wasps, but only against the hosts parasitised by *N. oneida.* We further investigated the pairwise differences in the cuticular hydrocarbon (CHC) profiles of the parasitised hosts and adult female wasps. The results reveal that the males show this ability only when the compounds responsible for differences in adult female CHC profiles were also the key differentiators of the host CHC profiles. The comparative mate searching behaviour of males of all reported species within a genus has rarely been studied. Therefore, this study makes a significant contribution to our understanding of interspecific variation of conspecific-mate searching behaviour.

## 1. Introduction

Successful reproduction requires the timely recognition of conspecific mates. Therefore, finely tuned mate recognition systems have evolved for both sexes in most species (reviewed in Wiley, 2013; Adams & Tsutsui, 2020). Additionally, the reproductive success of males is generally limited by the number of fertile females and the number of eggs laid by them (Bateman, 1948). Thus, sex-specific and divergent selective pressures can lead to the development of distinctive reproductive strategies that maximise fitness (Gross, 1996). Males can accomplish this directly by intense mate searching (reviewed in Parker, 1978), multiple mating (Bateman, 1948; Dewsbury, 2005), producing pheromones to attract females (Landolt, 1997), and attaining protandry or achieving reproductive maturity before the females (Andersson & Iwasa, 1996). Males can also employ several indirect strategies, like male-male competition (Andersson, 1994; Baer, 2020), territoriality (reviewed in Fitzpatrick & Wellington, 1983), monopolising access to females (reviewed in Parker, 1978), sneaky copulation (Küpper, 2021), etc. However, before employing any of these direct or indirect strategies, males must actively search for potential mates by using sensory cues (reviewed in Bell, 1990) to gain access to as many mates as possible (reviewed in Gröning & Hochkirch, 2008). When females are present in patches, males must also optimise the time and effort spent on each such patch (reviewed in Bell, 1990; Louâpre et al., 2015). Selection pressure for narrower conspecific mate recognition cues, however, can further intensify if a pool of possible mates contains closely related sympatric species, as speciation is not always accompanied by divergence in species recognition cues (reviewed in Gröning & Hochkirch, 2008; Mendelson & Shaw, 2012). This has been shown in many recently diverged taxa where species recognition cues are not distinctive and species-specific (Coyne et al., 2002; Gow et al., 2006; Gröning & Hochkirch, 2008; Mendelson & Shaw, 2012).

The parasitoid wasp *Nasonia* (Hymenoptera: Pteromalidae) is one of the most well-characterised insect systems for understanding sexual communication and is an excellent model system for uncovering conspecific mate-finding strategies (van den Assem, 1996; Drapeau & Werren, 1999; van den Assem & Beukeboom, 2004; Steiner et al., 2006; Ruther et al., 2007; Mair & Ruther, 2019; Prazapati et al., 2022). *Nasonia* is a parasitoid of cyclorrhaphous fly pupae (Whiting, 1967) and completes its development inside the host. The protandrous males usually wait on the natal host patch for females to emerge. Females, upon emerging, usually mate only once and fly out to seek fresh pupae to lay eggs (Whiting, 1967) by drilling a hole with their ovipositor (Martinson et al., 2014).

Among the four *Nasonia* species, *N. vitripennis* is cosmopolitan and therefore, sympatric with the other three species-*N. longicornis* (found in western North America), *N*. *giraulti*, and *N. oneida* (both found in eastern North America and micro-sympatric with each other across their range (Raychoudhury et al., 2010)). This sympatric distribution can affect the parasitisation of a patch of fly pupae by causing multi-parasitism, where the patch can be parasitised by multiple species of *Nasonia* (Wylie, 1965; Darling & Werren, 1990; Grillenberger et al., 2009). The reproductive interference caused by such multi-parasitism (reviewed in Noor, 1999; Gröning & Hochkirch, 2008) can be a selection pressure for the evolution of prezygotic barriers. Therefore, *Nasonia* males should be under selection to evolve precise mate recognition systems (van den Assem & Werren, 1994; Noor, 1999; Gröning & Hochkirch, 2008; Kyogoku, 2020). Evidence indicates that such a sympatric distribution may have given rise to some behavioural phenotypes in *Nasonia,* like the species-specific male courtship behaviour (van den Assem & Werren, 1994) and the evolution of within-host mating, particularly in *N. giraulti* (Drapeau & Werren, 1999; Trienens et al., 2021). Additionally, the pronounced female-biased sex ratio (Whiting, 1967) and avoidance of multiple mating by females (Van den Assem & Jachmann, 1999) can also intensify selection on males to develop precise species-recognition cues. This strategy can include the ability of males to detect hosts with conspecific females and thereby monopolizing access to emerging females. Accordingly, Prazapati et al. (2022) showed that male *Nasonia* can distinguish parasitised hosts from unparasitised ones, and *N. vitripennis* males can even detect which fly hosts have adult females inside them. This ability was found to be correlated with males’ ability to perceive female-specific cuticular hydrocarbons (CHCs) from the hosts (Prazapati et al., 2022). However, this strategy would only work to maximise male fitness if they can detect parasitised hosts with conspecifics. As shown above, theoretically, multiple selection forces should enable *Nasonia* males to distinguish hosts with conspecific mating partners. However, whether such abilities exist has not been studied empirically.

Here, we investigated if *Nasonia* males exhibit any such ability to detect hosts with conspecific wasps by providing them with a choice of hosts with conspecific and heterospecific wasps in a cafeteria assay. Further, we studied the searching behaviour of males by developing a customised Python programme for tracking the movement of males and looked for species-specific differences. Finally, we explored the CHC profiles of parasitised hosts and adult females to test the presence of species-specific olfactory signatures that can facilitate the searching behaviour of males.

## 2. Materials and methods

### 2.1 Strains used

We conducted all behavioural and CHC profile studies with four *Nasonia* species. The strains used were NV-IPU08 of *N. vitripennis*, NL-MN8501 of *N. longicornis*, NG-RV2XU of *N. giraulti,* and NO-NY11/36 of *N. oneida*. All strains were maintained at 24-hour constant light, at 25°C temperature with 60% relative humidity. The life cycles of the four species under these conditions were 14 days for *N. vitripennis*, 14.5 days for *N. longicornis*, 15 days for *N. giraulti*, and 16 days for *N. oneida*. We obtained virgin males and females by hosting virgin and mated females, respectively, on *Sarcophaga dux* (Diptera: Sarcophagidae) pupae. We obtained parasitised hosts for the cafeteria assay by exposing two hosts (less than 48h old) to a single mated female in a RIA vial for 48 hours. To ensure uniformity and minimise potential variability due to age and prior experience in the behavioural observations, we used virgin males, who were less than 24h old, and mated and virgin females, who were less than 48h old.

### 2.2 Cafeteria assay

We conducted the behavioural assays using an equal choice arena (Figure 1a), made of a glass Petri plate (9cm diameter), placed on a circular white paper of the same dimension. We drew two concentric circles of 9cm and 5cm diameters on the paper and divided the annulus into six equal sections (Prazapati et al., 2022). We alternatively placed 6 hosts (3 containing conspecific and 3 with heterospecific wasps) on these six sections. This setup was placed on a wooden platform with a white LED lamp above at a height of 30 cm. To restrict the males within the arena, we made a thin water ring at the circumference of the Petri plate. We then introduced a virgin male in the centre of this arena and recorded its behaviour for the next 4 minutes using a Logitech C615 HD camera (all data are uploaded at https://www.youtube.com/@Evogeniiserm). We intended to use hosts with mature wasps about to eclose. Therefore, to confirm the parasitisation of the hosts with the required pupal stage, we opened each host pupa after the behavioural assay and verified the developmental stage. We discarded the assays where one or more hosts failed to meet this criterion. The males of each species were subjected to 3 categories of assays, where their behaviour was recorded for 4 minutes. This resulted in 12 different categories of assays across the 4 species, with 30 data points for each category. Each data point was collected using a fresh set of Petri plates, hosts, and inexperienced virgin males.

**Figure 1:**
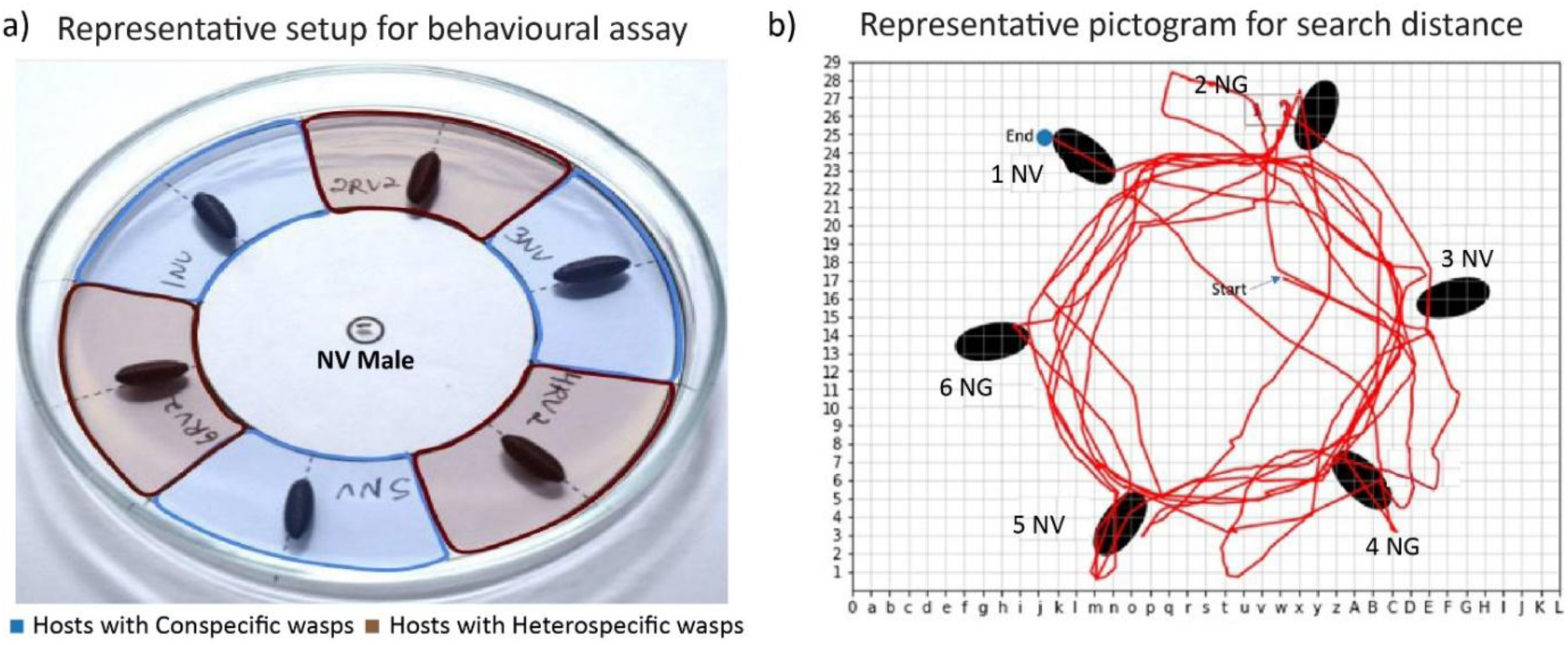
a) Cafeteria Arena set up for the behavioural assays: 3 hosts parasitised by conspecific females and 3 hosts parasitised by heterospecific females were placed alternatively. In each assay, one male was introduced at the centre of the plate. Video recording was done immediately after introducing the male for 4 minutes using a Logitech C615 HD webcam. The colours used to distinguish hosts in the picture are only for representation. b) The distance traversed by the males: Movement of each male was tracked and digitally scored by the customised Python program for measuring Search Distance and calculating Search Speed.

To assess whether males can identify hosts with conspecific wasps, we compared the total time each male spent on the two kinds of hosts. The times spent on hosts with conspecific or heterospecific wasps were calculated by summing the total time spent on all three hosts of similar type. Time spent was recorded from the moment a male climbed on a host and ended when it descended from it. If the males spent significantly more time on hosts with conspecific wasps, we considered them capable of preferring or identifying the hosts with conspecific over hosts with heterospecific wasps.

### 2.3 Searching Behaviour

To quantify the different parameters of ‘searching behaviour,’ we designed a customised Python program, which tracked each male’s movement in the arena for the entire duration of the behavioural assay (Figure 1b, SI Python script). We incorporated four possible search parameters (Table 1) to assess variation in search behaviour: 1) ‘search distance’, 2) ‘search time’, 3) ‘search speed’, and 4) ‘search latency’. For this analysis, we used the 24 videos from each category of bioassay and pooled the data for males of a given species across the three choice assays.

**Table 1:** Defines the list of search parameters included in the study to analyse the search behaviour of males and their quantifications.

| Search parameters | Definition | Quantification |
| --- | --- | --- |
| Search distance | Total distance (in meters) travelled by individual males in the arena for 4 minutes. | Calculated using a customised program written in Python (SI Python Script, Figure 1b), which tracked the movement of males. |
| Search time | The total duration of time for which males were moving around and not stationed on a particular host. | Calculated by subtracting the time spent on hosts from the total assay time (4 minutes). |
| Search speed | The average speed at which males were moving in the cafeteria arena. | Calculated by dividing the search distance by the search time in m/sec. |
| Search latency | The time taken by a male (in seconds) to find the first host with conspecifics. Finding was scored when a male climbed on a host. | Calculated as the time from the start of the assay till the male found the first host with conspecifics. |

### 2.3 Purification and identification of Cuticular Hydrocarbons (CHCs)

We analysed the CHCs of the parasitised hosts and the adult virgin females of all four species. For these eight categories, we had three replicates, each prepared from 50 individual hosts or 50 females. We collected CHCs from parasitised hosts (13 to 15 days old, depending on the *Nasonia* species) by dipping them in 3 ml of n-hexane (HPLC grade MERCK) for 5 min. The acquired solution was then concentrated to 100 µl under a gentle nitrogen stream. The process was repeated for adult females, which were similarly dipped in 500 µl n-hexane for 10 min. The collected extracts were then processed to concentrate the CHCs using column chromatography in glass columns (0.7 cm inner diameter), which were packed 3cm high with baked glass wool and silica gel. Eluted fractions of hosts and females were then separately concentrated to a 50 μl volume in a gentle nitrogen stream.

To identify the chemical signature of these extracts, the samples were run through a gas chromatograph connected with a mass spectrometer (Agilent 7890B, 5977C GC-MS) with an HP-5MS (Agilent J&W) capillary column (30 m X 0.25 mm X 0.25 μm), operated in a 70-eV electron impact ionization mode. The quadrupole temperature was maintained at 150°C, and the inlet and auxiliary line temperatures were kept at 320°C. 2 μl (1 host or female equivalent/μl) of the extracted sample was injected individually in a split-less mode into the GC machine, where Helium was used as the carrier gas at an average linear velocity of 36.2 cm s^-1^. Separation of these fractions was advanced, keeping the initial temperature at 40°C, which was held for 5 minutes. The oven temperature was then raised to 320 °C.

The percentage of the relative abundances was calculated for each detected compound (Tables S9 and S10). The identification of the compounds involved a comparison of the distinctive diagnostic ions present in the obtained mass spectra with the NIST library of Mass Hunter Workstation Software vB.08.00 (Agilent Technologies) and *n*-alkane standards (*n*-C8 to *n*-C40; SIGMA-40147-U) (Prazapati et al., 2022). Methyl-branched hydrocarbons were identified through their characteristic diagnostic ions arising from the fragmentations at their branching points and the utilisation of linear retention index (LRI) values from previously published data (Carlson et al., 1999; Steiner et al., 2006; Prazapati et al., 2022).

To identify the potential CHC compounds responsible for host recognition by the males, we parsed the CHC profiles from hosts parasitised by each species of females. The full GC profiles and their corresponding SIMPER analyses are given in Tables S11-S15. We focused on the unique peaks as well as peaks with varying relative abundances (Soon et al., 2021), which together contributed to 50% of the total dissimilarity between these profiles (Jungwirth et al., 2021). This was also done for the adult female profiles (the full GC profiles and their corresponding SIMPER analyses are given in Tables S16-S20), as hosts contain more females than males due to the female-biased sex ratio of *Nasonia* (Whiting, 1967).

## 3. Statistical analyses

We assessed the data distribution using the Shapiro-Wilk’s test for Normality (Shapiro & Wilk, 1965) and used parametric and non-parametric tests for normally distributed data and non-normally distributed data, respectively. We employed the Wilcoxon signed-rank test for the time spent on hosts by males. We used one-way ANOVA, followed by Tukey’s HSD *post hoc* test, to compare interspecific differences in search time and speed. In contrast, for analysis of the other two search parameters, search distance and search latency, we used the Kruskal-Wallis test, followed by Dunn’s *post hoc* test. We conducted Principal components analysis (PCA) on the CHC profiles. Prior to carrying out the PCA, we added a small constant to the dataset before transformation to address the presence of undetectable compounds in some samples (Hervé et al., 2018) and then transformed the data using centred log-ratio (CLR) (Aitchison, 1982). We identified the CHCs responsible for contributing to species-specific differences by a Similarity Percentage (SIMPER) analysis, which ranks the contribution of individual CHC peaks based on relative abundance and presence and/or absence of the compounds by Bray-Curtis distance measures (CLARKE, 1993) using Past4.03.exe (Hammer & Harper, 2001). We used R v. 4.2.2 (R Core Team, 2024) for data transformation and Origin 2022 (OriginLab Corp.) for all statistical analysis and constructing the box plots and bar graphs.

## 4. Results

### 4.1 *N. vitripennis* and *N. longicornis* males can identify hosts with conspecific wasps

We found a distinct pattern of host preference in the choice assays where males of all four *Nasonia* species were placed with hosts with conspecific and heterospecific wasps (Figure 2, Tables S1-S4). Males of *N. vitripennis* spent significantly more time on the host with conspecifics against *N. giraulti* and *N. oneida* (Wilcoxon signed-rank test, N = 30, *p* = 0.005 and N = 30, *p* = 0.001, respectively; Figure 2a). However, they did not show any preference for hosts with conspecifics against hosts parasitised by *N. longicornis* and spent nearly equal time on both (Wilcoxon signed-rank test, N = 30, *p* = 0.265; Figure 2a). This indicates that *N. vitripennis* males can identify hosts with conspecifics from hosts with heterospecifics, except when the latter is parasitised by *N. longicornis*. Males of *N. longicornis* spent significantly more time on hosts with conspecifics only against *N. oneida* (Wilcoxon signed-rank test, N = 30, *p* = 0.002; Figure 2b), but did not show any preference to hosts with conspecifics against hosts parasitised by *N. vitripennis* and *N. giraulti* (Wilcoxon signed-rank test, N = 30, *p* = 0.918 and N = 30, *p* = 0.464, respectively; Figure 2b). Males of *N. giraulti* did not exhibit any preference in time spent on hosts with conspecifics over hosts parasitised by *N. vitripennis*, *N. oneida,* and *N. longicornis* females (Wilcoxon signed-rank test, N = 30, *p* = 1; N = 30, *p* = 0.569 and N = 30, *p* = 0.127, respectively; Figure 2c). Similarly, males of *N. oneida* failed to distinguish between hosts with conspecific and those parasitised by *N. giraulti* and *N. longicornis* (Wilcoxon signed-rank test, N = 30, *p* = 0.077 and N = 30, *p* = 0.589, respectively; Figure 2d). Interestingly, *N. oneida* males spent significantly more time on hosts parasitised by *N. vitripennis* females (Wilcoxon signed-rank test, N = 30, *p* = 0.037; Figure 2d) than on hosts parasitised by conspecific females.

**Figure 2:**
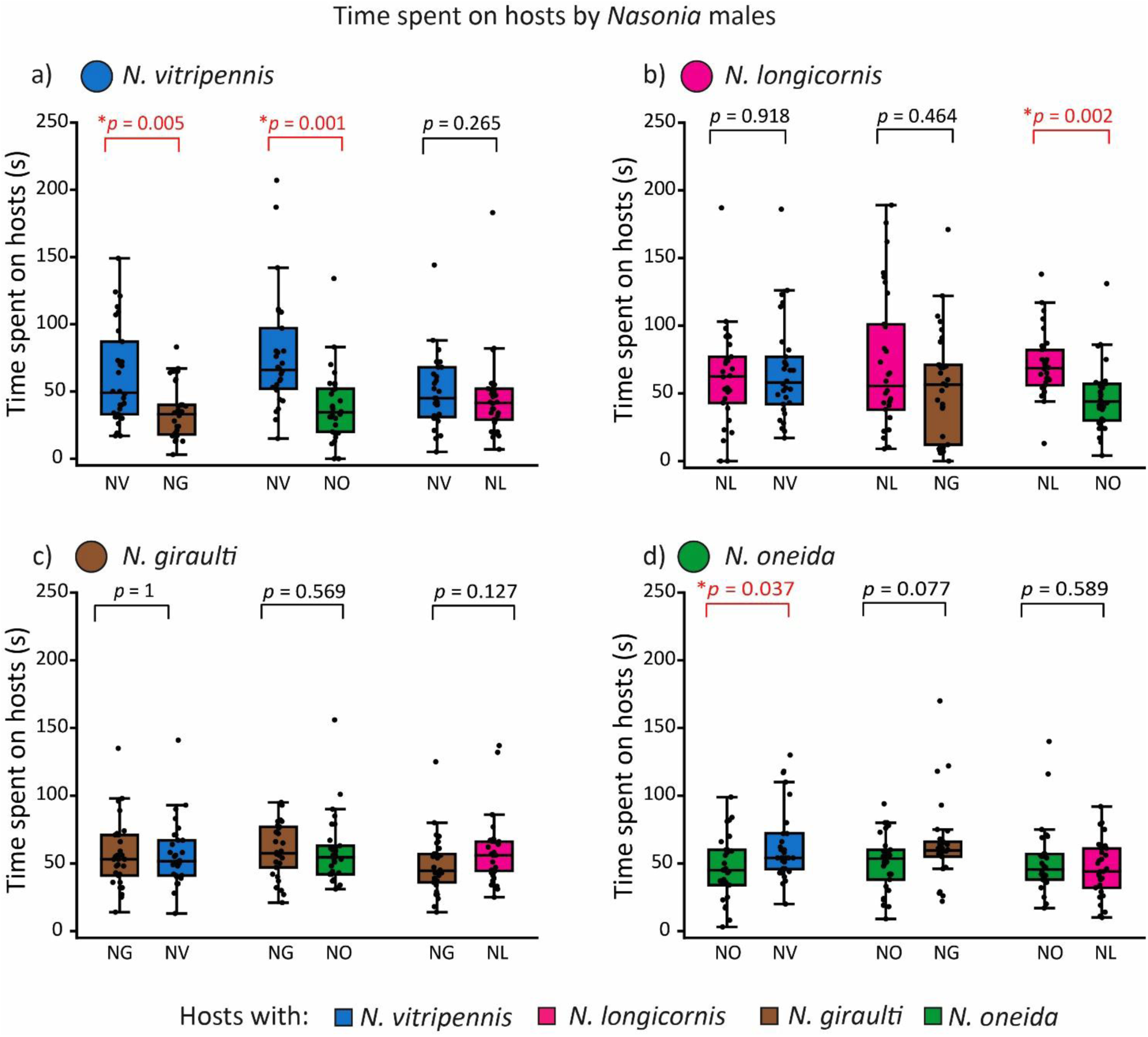
Comparison of wait duration by *Nasonia* males on hosts with conspecific wasps and hosts with heterospecific wasps in the cafeteria assay: a) *N. vitripennis* males spent significantly more time on hosts with conspecific against hosts with *N. giraulti* and *N. oneida* b) *N. longicornis* males spent significantly more time on hosts with conspecific against hosts with *N. oneida* c) *N. giraulti* males did not spend significantly longer duration on any host d) *N. oneida* males spent significantly more time on hosts with *N. vitripennis* but spent similar time on both kinds of hosts against *N. giraulti* and *N. oneida.* Each comparison is based on data for 30 assays, and asterisks represent significant differences (Wilcoxon signed-rank test, *p* < 0.05).

### 4.2 *N. vitripennis* and *N. longicornis* males use different strategies while searching

As the above results indicate, *Nasonia* males have varying abilities to identify hosts with conspecifics. To understand whether this varying ability is an outcome of how these males search for conspecifics, we compared the search behaviour of four species by exploring four parameters: search distance, search time, search speed, and latency to find the first host with conspecifics (search latency) (Table 1).

We compared the distance traversed (search distance) by males in 4mins, and found that *N*. *vitripennis* males cover the longest distance, which was significantly more than males of other three species (Kruskal-Wallis with Dunn’s test N =72, *p* = 0.004, N = 72, *p* = 0.003, N = 72, *p* < 0.0001 with *N. giraulti*, *N. oneida*, *N. longicornis* respectively; Figure 3a, Table S5). There was no significant difference in the distances covered by *N. giraulti* and *N. oneida* (Kruskal-Wallis with Dunn’s test, N = 72, *p* > 0.05; Figure 3a, Table S5), while *N. longicornis* covered the least distance (Kruskal-Wallis with Dunn’s test, N = 72, *p* = 0.001, N = 72, *p =* 0.001, for *N. giraulti* and *N. oneida,* respectively; Figure 3a, Table S5). Therefore, the success of *N. vitripennis* in being the most successful species in identifying conspecific mates could be due to the effort that each male spends in covering the longest search distance.

**Figure 3:**
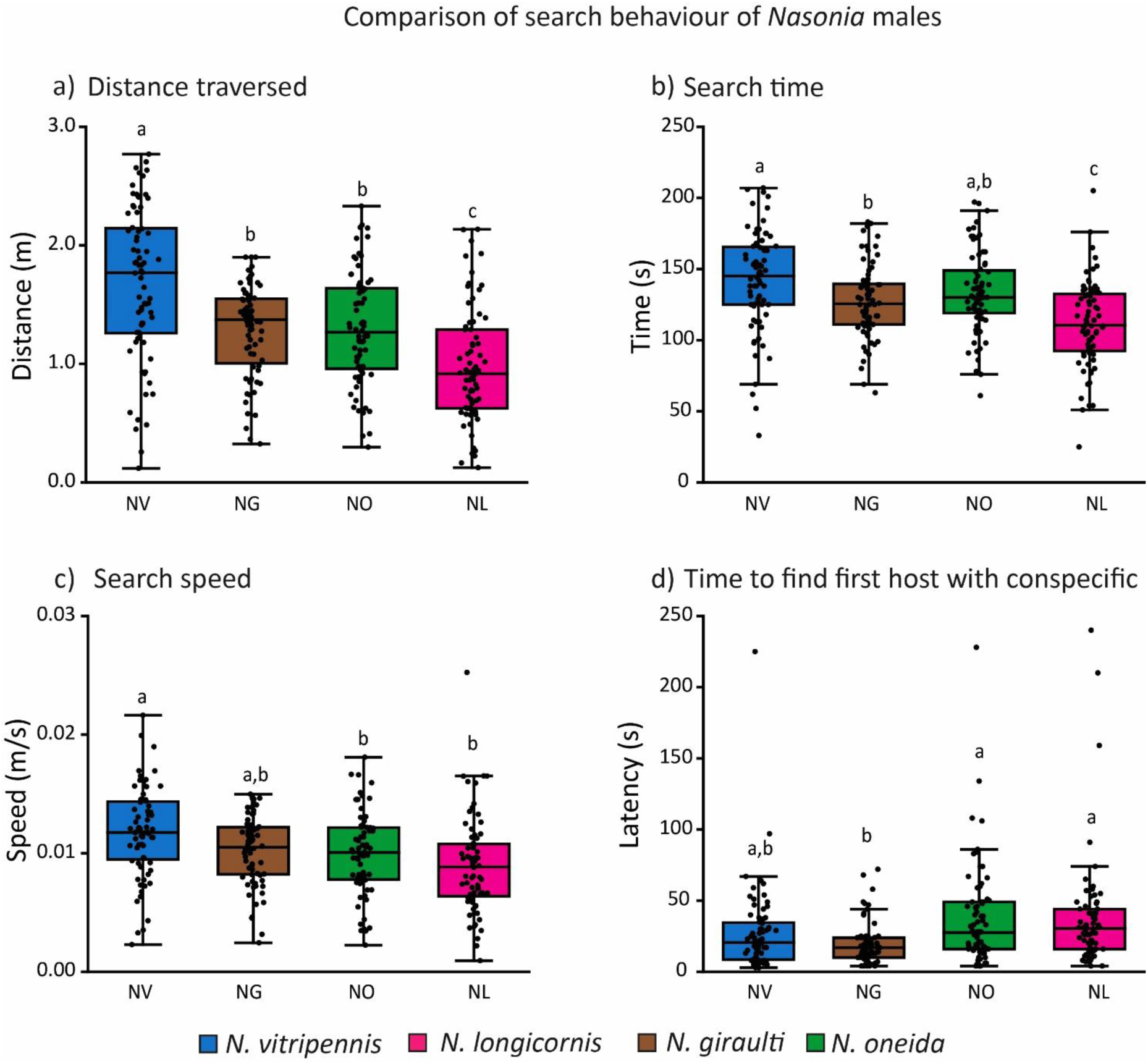
Comparison of 4 parameters of search behaviour of *Nasonia* males: a) Search Distance, b) Search Time, c) Search Speed, and d) Search Latency. Boxes carrying different alphabets represent significant differences. (Kruskal-Wallis with Dunn’s as post hoc test, *p* < 0.05 for Search Distance and Latency, and one-way ANOVA with Tukey’s HSD *post hoc* test, *p* < 0.05 for Search Time and Search Speed. For each species, 24 data points were used, and based on the data distribution, parametric or non-parametric tests were employed.

As we compared the search time of the four species, we found that *N. vitripennis* males also spent the most time searching for hosts, which was significantly longer than *N. giraulti* but not significantly different from *N. oneida* males (one-way ANOVA with Tukey’s HSD, N = 72, *p* = 0.027, N = 72, *p* = 0.364, for *N. giraulti* and *N. oneida* respectively; Figure 3b, Table S6). There was no significant difference in search time between *N. giraulti* and *N. oneida* males (one-way ANOVA with Tukey’s HSD, N =72, *p* = 0.63705; Figure 3b, Table S6). However, the search time for all three species was significantly longer than that of *N. longicornis* males (one-way ANOVA with Tukey’s HSD, N = 72, *p* < 0.001, N = 72, *p* = 0.010, N = 72, *p* < 0.001, *N. vitripennis*, *N. giraulti*, *N. oneida*, respectively; Figure 3b, Table S6).

We further compared the search speed (a function of the above two parameters) of the males. Males of *N. vitripennis* exhibited the fastest search speed (Table S7), which was significantly different from that of *N. longicornis* and *N. oneida* males but not from *N. giraulti* males (One-way ANOVA with Tukey’s HSD, N =72, *p* < 0.001, N = 72, *p* = 0.013, and N = 72, *p* = 0.056 for *N. longicornis*, *N. oneida* and *N. giraulti,* respectively; Figure 3c). However, the search speed of the males of the other three species was not significantly different from each other (one-way ANOVA with Tukey’s HSD, N = 72, *p* > 0.05; Figure 3c, Table S7).

Although *N. vitripennis* males significantly differed from the other species in the above three parameters, they did not differ in the latency to find the first host with conspecific wasps (Kruskal-Wallis with Dunn’s test, N = 72, *p* > 0.05; Figure 3d). *N. giraulti* exhibited a shorter latency than *N. longicornis* and *N. oneida* (Kruskal-Wallis with Dunn’s test, N = 72, *p* < 0.001, N = 72, *p < 0.001* for *N. longicornis* and *N. oneida,* respectively; Figure 3d, Table S8), but it was not significantly different from *N. vitripennis* (Kruskal-Wallis with Dunn’s test, N = 72, *p* = 0.390; Figure 3d, Table S8).

### 4.3 Comparative analysis of olfactory cues

The GC-MS analysis of the CHC profile of the parasitised hosts (having adults inside) and adult females shows the presence of long-chain saturated and unsaturated hydrocarbons ranging from nC25 to nC37 (Figure 4, Table S9 and S10, respectively), with the majority composed of methyl-branched alkanes. 46 peaks were identified in host CHC profiles, out of which 43, 34, 33, and 38 peaks were found in hosts parasitised by *N. vitripennis*, *N. giraulti*, N*. oneida*, and *N. longicornis* females, respectively (Table S9). In contrast, 50 peaks were identified in CHC profiles of adult females with 4 additional peaks: MeC25(5-), MeC27(7-), MeC27(5-), and MeC28(4-). Specifically, 48, 48, 29, and 45 peaks were found in *N. vitripennis*, *N. giraulti*, *N. oneida*, and *N. longicornis* females, respectively (Table S10). The identified peaks in the female CHC profile contain both previously reported (Buellesbach et al., 2013) and newly identified peaks.

**Figure 4:**
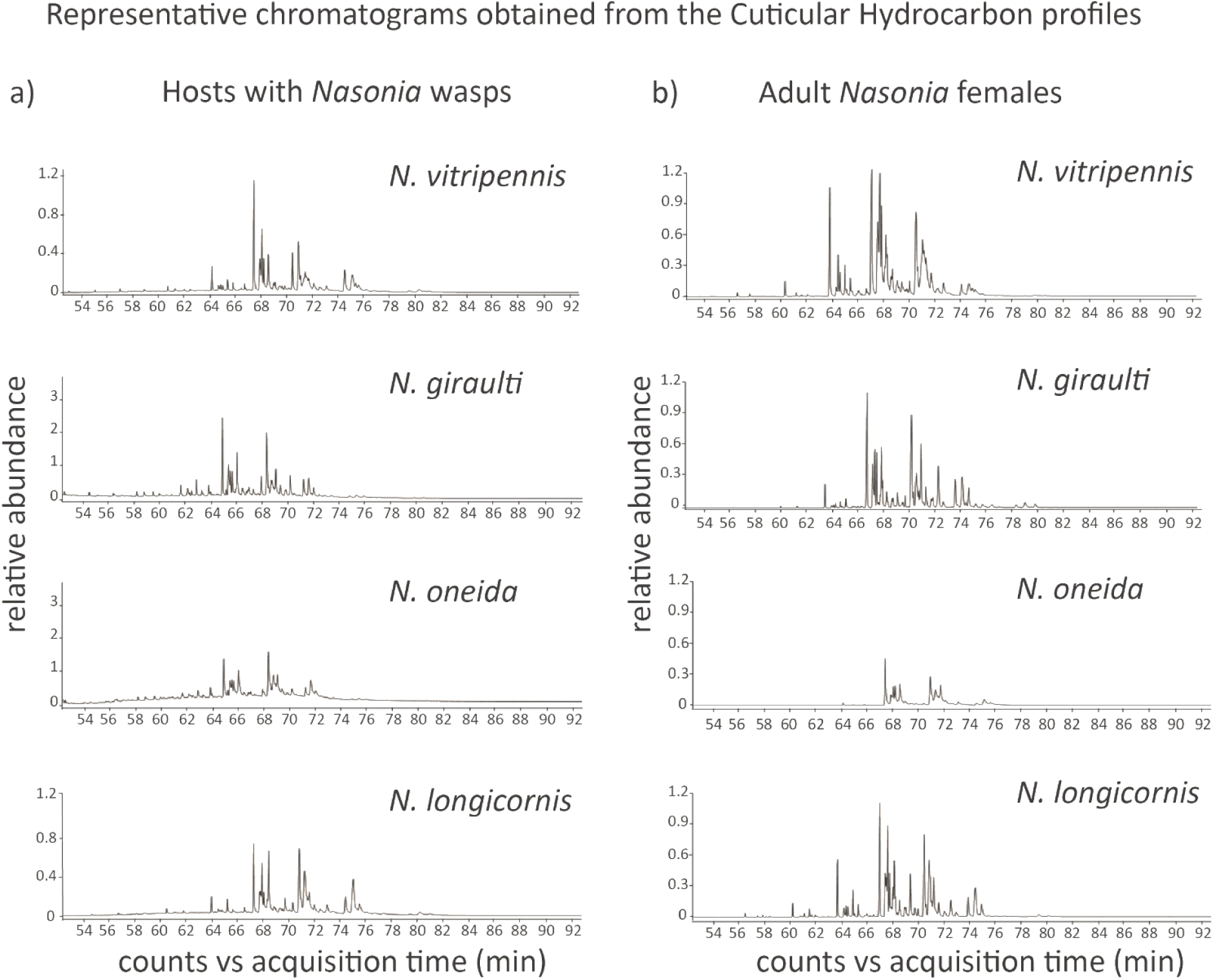
a) Representative chromatograms obtained from the CHC profiles of hosts with adult males and females inside. A total of 46 peaks were identified in CHC profiles of hosts parasitised by *Nasonia* females of different species. b) Representative chromatograms obtained from the CHC profiles of adult *Nasonia* females. A total of 50 peaks were identified in CHC profiles of *Nasonia* females of different species.

*Nasonia* males can utilise olfactory cues in the form of female CHCs to distinguish different types of hosts (Prazapati et al., 2022). In the choice assays of this study, we found that *N. vitripennis* males can detect the hosts with conspecific wasps against hosts with *N. giraulti* and *N. oneida*, and *N. longicornis* males can achieve this against hosts with *N. oneida.* To understand the potential olfactory cues responsible for the identification, we compared the CHC profiles of every combination of hosts offered in the choice assays to these two species. Further, to investigate if the host-CHCs had any similarity with the CHCs of the females within, we further compared the CHC combinations of females.

7 different peaks contributed to 50% of the dissimilarity observed between the hosts parasitised by *N. vitripennis* and *N. giraulti* (Figure 5a, Table S11). Out of these, 3 peaks [C31, MeC33(3-) and MeC33(15-;13-;11-)] were also found to contribute to 50% of the dissimilarity seen in the adult female profiles of these two species (Table S16). Similarly, the comparison between hosts parasitised by *N. vitripennis* and *N. oneida* also yielded 7 such peaks (Figure 5b, Table S12). Out of these, 4 peaks [C31, MeC33(3-), MeC31(7-), and MeC33(5-)] contributed to 50% dissimilarity of adult females of these two species (Table S17). These results suggest that in cases of successful identification, the main compounds responsible for the dissimilarity in females can also contribute to the olfactory uniqueness of the hosts. This contention is further strengthened by the comparisons with indistinguishable hosts parasitised by *N. vitripennis* and *N. longicornis,* where 6 peaks contributed to 50% dissimilarity (Figure S1a, Table S13), but only one of them, MeC35(15-;13-;11-), was common to those contributing to 50% dissimilarity between the females of corresponding species (Table S18).

**Figure 5:**
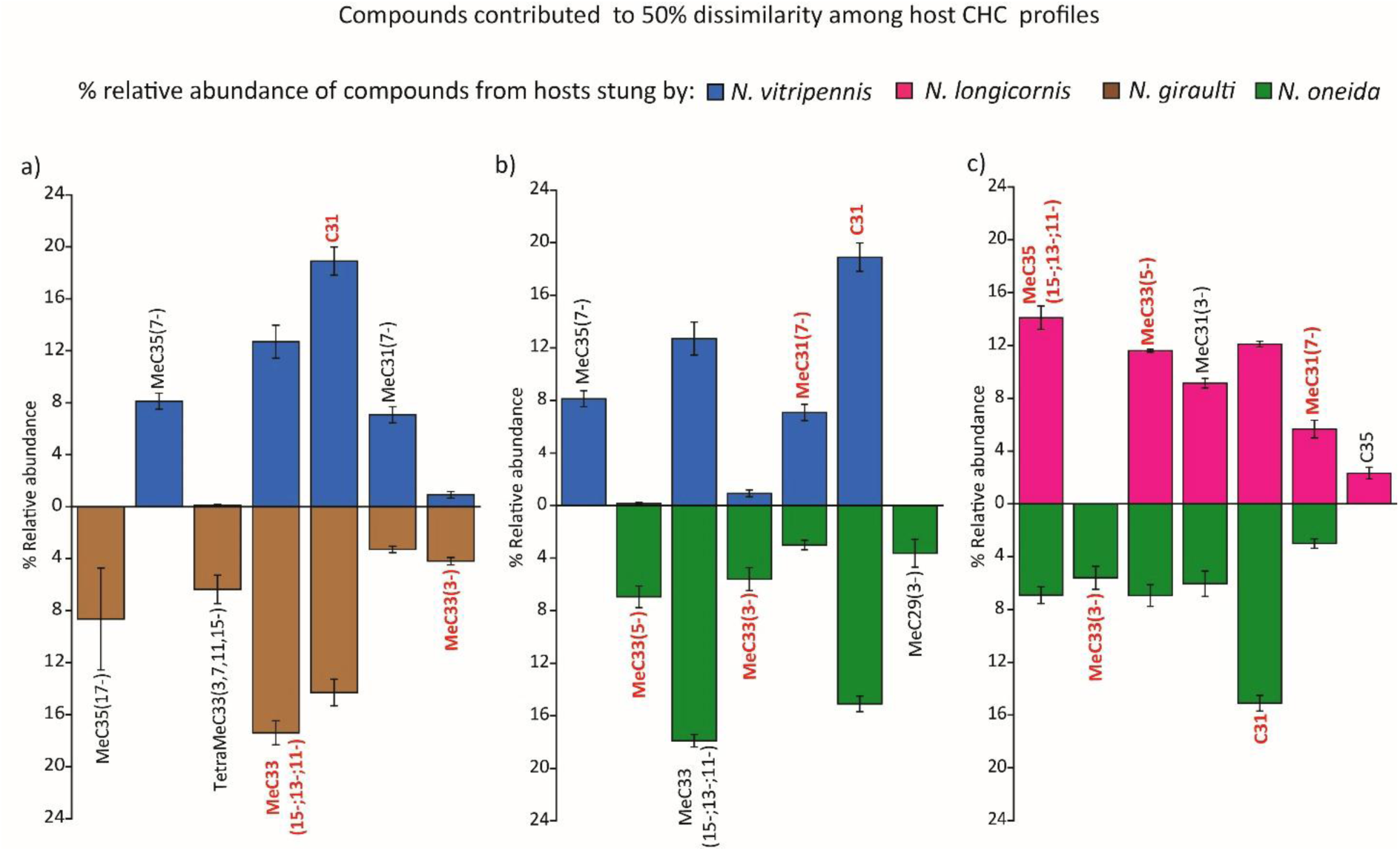
Relative abundance (%) of compounds that cumulatively contributed to 50% of dissimilarity between CHC profiles of parasitised hosts, which were distinguished by males: Comparison of compounds between a) hosts parasitised by *N. vitripennis* and *N. giraulti,* b) *N. vitripennis* and *N. oneida,* c) hosts parasitised by *N. longicornis* and *N. oneida*. Compounds titled in red also contribute to dissimilarity in adult female CHC profiles of the corresponding species.

*N. longicornis* males were able to identify the hosts with conspecific wasps only against hosts parasitised by *N. oneida.* The CHC comparison of hosts parasitised by these two species revealed 7 peaks contributing to 50% dissimilarity of the hosts (Figure 5c, Table S14). Among them, 5 peaks [C31, MeC33(3-), MeC33(5-), MeC31(7-), and MeC35(15-;13-;11-)], were also found to be different between the adult females of these two species (Table S19). Comparison between hosts parasitised by *N. longicornis* and *N. giraulti* showed 5 peaks (SI Figure S6b, SIMPER analyses SI Table S115) to be different, but shared only one, MeC33(3-), with the adult female profiles (Table S20).

These results indicate that successful identification by *N. vitripennis* and *N. longicornis* males may require the presence of a blend of female-specific CHCs in the host, as when these peaks were absent, the males failed to identify hosts with conspecifics.

The above results aligned with the Principal Components analysis (PCA) of the relative abundance (%) for the CHCs extracted from the parasitised hosts and females of four species (Figure 6). The first two principal components explained 56.46% variation, and when plotted against each other, separations were seen among the CHC profiles of parasitised hosts and females (Figure 6). The hosts parasitised by *N. vitripennis* and *N. longicornis* were clustered together and away from the cluster of hosts parasitised by *N. giraulti* and *N. oneida.* Furthermore, the profiles of hosts parasitised by *N. vitripennis* and *N. longicornis* were close to the profiles of their respective females, but the hosts parasitised by *N. giraulti* and *N. oneida* were relatively far from their respective females.

**Figure 6:**
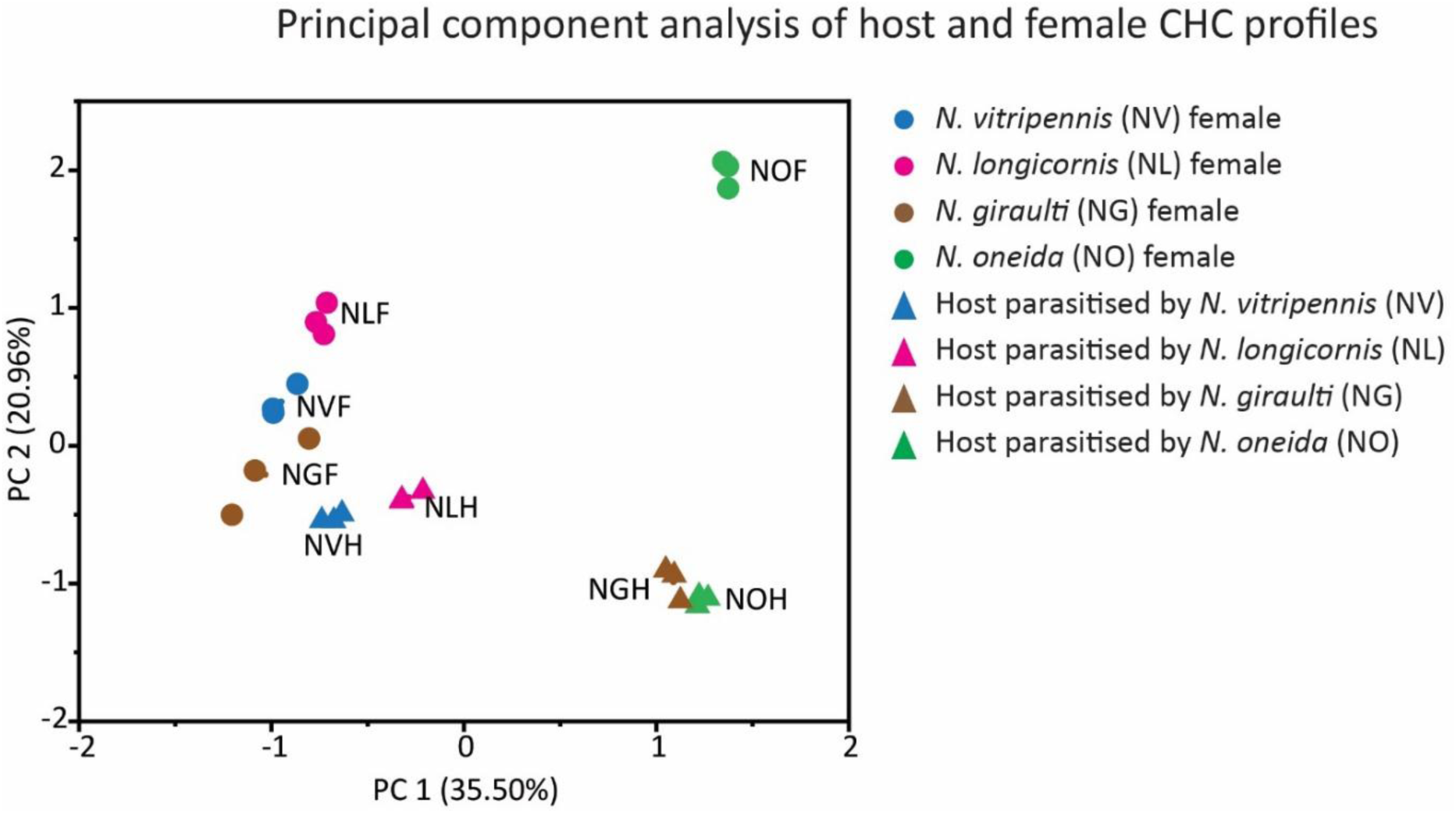
Distribution of CHC profiles of *Nasonia* females (circles) and parasitised hosts (triangles) in two-dimensional principal components space.

## 5. Discussion

*Nasonia* males can mate with numerous females in quick succession, limited only by the availability of females (Tiwary et al., 2022). The females emerge from hosts (fly pupae), which usually occur in batches, and can be parasitised by more than one species of *Nasonia* (Wylie, 1965; Darling & Werren, 1990; Grillenberger et al., 2009). Therefore, to maximise the number of successful matings, a male must find hosts that are parasitised by conspecific wasps and wait for the females to emerge. Moreover, the emerging *Nasonia* females mate only once before flying away to look for fresh hosts to parasitise (Whiting, 1967; King, 1993; King et al., 2000). Therefore, the males should be under selection to optimise their energy expenditure and search time to locate hosts with conspecific females and wait. We expected that (i) males should be able to identify the hosts with conspecific wasps and exhibit that by spending more time on them, and (ii) they should optimise the duration of waiting on a host and energy expenditure in terms of distance traversed and speed of search. To test these hypotheses, we placed three hosts with conspecific wasps and three hosts with heterospecific wasps in a cafeteria assay. We considered a longer wait (time spent) on the hosts with conspecific wasps to be a measure of the males’ ability to identify the hosts with conspecific wasps. We observed that *Nasonia* males could identify hosts with conspecific wasps, but this ability was not uniform across the species or even within each species. *N. vitripennis* showed this ability against *N. oneida* and *N. giraulti* but not against *N. longicornis*. Whereas, *N. longicornis* showed this ability only against *N. oneida*. *N. giraulti* and *N. oneida* did not exhibit any recognition of hosts with conspecifics (Figure 2). Moreover, our results also indicated that among the two species that had this capability, the pattern of searching was noticeably different (Figure 3). *N. vitripennis* searched for longer durations, travelling the longest distance and at the quickest speed, while *N. longicornis* males traversed the shortest distance with the slowest speed and searched for shorter durations than *N. vitripennis*. These results indicate that recognition of hosts with conspecific wasps can be achieved by both strategies in terms of energy expenditure.

To corroborate the role of olfactory cues in the asymmetry of recognition ability of the four species of *Nasonia*, we explored the CHC profiles of the parasitised hosts and females of *Nasonia*. It is well established that, except for *N. giraulti, Nasonia* males can identify and respond to female-derived CHCs (Steiner et al., 2006; Buellesbach et al., 2013; Mair et al., 2017; Sun et al., 2023). It has also been suggested that males likely perceive the cues left by females while ovipositing and/or emanating from the adult females developing inside the host for identification (Prazapati et al., 2022). The porous structure of the fly puparium (Yoder & Denlinger, 1991) can also facilitate males in perceiving the olfactory cues from females within. The volatility of the CHC compounds depends on carbon chain lengths, which is inversely proportional to the degree of volatility (Drijfhout et al., 2009). The compounds present on both the hosts and the females had long carbon chains (Tables S9-S10), which indicated low volatility. Therefore, we conjecture that the cumulative cues from the low volatility compounds within the host may provide a perceivable signal for the males. Our results suggest that successful identification corresponds to the compounds that contribute to the maximum difference in CHC profiles of hosts, which are also present in the female profiles. This indicates that males probably identify hosts with conspecifics by identifying the cues derived from adult females still inside the host. The effectiveness of these low-volatile CHCs serving as the cue is also consistent with the males’ inability to fly and being confined to their natal patch. Further empirical validation by isolating and testing these compounds is required to understand the nature of CHC utilisation by males.

The asymmetric and species-specific nature of the recognition appears to result from the probability of encountering heterospecific species based on the geographic distribution of the four species. The sympatric distribution can lead to selection for enhanced species-specific ability to distinguish mates (Brown & Wilson, 1956; Coyne & Orr, 1989; Howard, 1993; Butlin, 1995; Buellesbach et al., 2014; Raychoudhury, 2015). Accordingly, *N. vitripennis*, being cosmopolitan and sympatric with the other three species (Raychoudhury et al., 2010) shows the most extensive ability. However, this biogeographic argument is inconsistent with its inability to distinguish the sympatric *N. longicornis*. Moreover, *N. vitripennis* and *N. longicornis* showed a reciprocal inability to distinguish each other’s hosts with conspecific wasps. The other possibility could be a strain-specific effect. The *N. vitripennis* strain used (NV-IPU08) in this study is an Indian strain, which probably does not encounter any heterospecifics, as there is no report of any other species of *Nasonia* from India. However, this cannot explain why *N. vitripennis* can distinguish conspecifics against hosts stung by *N. giraulti* and *N. oneida*. But future studies should use *N. vitripennis* field strains from western North America, which are truly sympatric with *N. longicornis*, to understand whether such sympatric distribution can be correlated with this ability. This biogeographic explanation is also consistent with *N. longicornis’* inability to distinguish hosts with conspecific wasps over hosts parasitised by the allopatric *N. giraulti.* However, it can distinguish hosts with conspecific wasps from hosts parasitised by *N. oneida*, which is largely distributed across eastern North America and is allopatric to *N. longicornis*. The distribution of *N. oneida,* although concentrated in Eastern North America, can extend to the Midwest till Lake Mendota (Wisconsin, USA, Raychoudhury and Werren, personal observation). This indicates that *N. longicornis* and *N. oneida* can encounter each other in their native ranges and, therefore, can develop species-specific cues for the identification of hosts with conspecific wasps.

*N. giraulti* shows very high rates of within-host mating (Drapeau & Werren, 1999; Trienens et al., 2021). Since female *Nasonia* usually mate only once (Van den Assem & Jachmann, 1999), there are substantially fewer opportunities to mate with *N. giraulti* females outside the host. This indicates *N. giraulti* males should not be under selection to identify hosts with conspecific wasps. Our results affirm this prediction. Conversely, the low incidence of within-host mating in *N. vitripennis* and *N. longicornis* (Drapeau & Werren, 1999; Trienens et al., 2021) can be a strong selection pressure to correctly identify mates, which can explain why these are the only two species that can distinguish hosts with conspecifics. Moreover, both these species can show territorial and aggressive behaviour towards other males (van den Assem, 1996; Leonard & Boake, 2006; Mair & Ruther, 2018), indicating that they can be territorial in guarding hosts with females within them. However, both the biogeographic and within-host mating arguments cannot explain why *N. oneida* does not show this capability, as it is sympatric to both *N. vitripennis* and *N. giraulti* (Raychoudhury et al., 2010), as well as exhibits intermediate rates of within-host mating (Trienens et al., 2021). One possibility could be the loss of this ability in the most recent common ancestor of *N. oneida* and *N. giraulti,* which remains to be verified. However, *N. oneida* females exhibit the strongest mate preference (Raychoudhury et al., 2010) for conspecifics amongst all the *Nasonia* species. This can result in reduced selection pressure on the males to identify hosts with conspecific wasps, as the females will rarely mate with any heterospecific males. But why *N. oneida* males would prefer hosts parasitised by *N. vitripennis* females (Figure 2d) is difficult to explain. Such paradoxical behaviour by *N. oneida* males has also been documented in a previous study (Buellesbach et al., 2014), where they reject conspecific females at nearly the same rate as heterospecific ones.

*Nasonia* is one of the best-characterised insect model systems to study sexual communication (van den Assem, 1996; Drapeau & Werren, 1999; van den Assem & Beukeboom, 2004; Steiner et al., 2006; Ruther et al., 2007; Mair & Ruther, 2019). However, as *Nasonia* is also a flagship genus for studying local mate competition and sex ratio variations, most studies have focused only on females (Werren, 1980, 1983; King, 1992; Shuker et al., 2006; Burton-Chellew et al., 2008). Recent studies (Leonard & Boake, 2006, 2008; Mair & Ruther, 2018; Kurtanovic et al., 2022; Prazapati et al., 2022) have also revealed the complexities of male behaviour. This study unveiled a much more varied and complex reproductive behavioural repertoire of *Nasonia* males than previously known.

## Statements and Declarations Ethics

The entire work with wasps was carried out following all the safety measures of the Institute Biosafety Committee of the Indian Institute of Science Education and Research, Mohali, India.

## Consent for Publication

All authors agree to the submission of the manuscript, and Rhitoban Raychoudhury has been authorized as the corresponding author by all co-authors.

## Availability of data and material

All behavioural assay videos are available at https://www.youtube.com/@Evogeniiserm. Python script used to track the movement of males, and other data is given in the supplementary information (Verma et al., 2025).

## Declaration of AI use

We have not used AI to create this article.

## Author contribution

TV and RR conceived and designed the study. TV collected and analysed data and developed the script with BKS. DB analysed the CHC profiles. SJ and AA provided resources. TV and RR wrote the first draft of the paper. RS reviewed and edited the manuscript.

## Acknowledgement

We thank Garima Prazapati for initially finding such behavioural variations and providing guidance.

