## Supplementary material for "*Nasonia vitripennis* males exhibit greater effort and competency in detecting hosts with conspecific females than other *Nasonia* males": Supplimentary Information (SI)

**This file includes**

**Table S1 to S20**

**Figure S1**

**Python Script: Code for tracking *Nasonia* male movement**

**Table S1:** Time spent by *N. vitripennis* males on hosts with conspecifics and hosts with heterospecifics in the cafeteria assay, having three different categories.

| S.no. | Male ID | Time spent on hosts by NV male (sec) | | Male ID | Time spent on hosts by NV male (sec) | | Male ID | Time spent on hosts by NV male (sec) | |
|---|---|---|---|---|---|---|---|---|---|
| | | Hosts with conspecifics (NV) | Hosts with heterospecifics (NG) | | Hosts with conspecifics (NV) | Hosts with heterospecifics (NO) | | Hosts with conspecifics (NV) | Hosts with heterospecifics (NL) |
| 1 | 15 | 124 | 58 | 53 | 53 | 19 | 6 | 28 | 20 |
| 2 | 24 | 73 | 24 | 59 | 63 | 43 | 14 | 56 | 32 |
| 3 | 34 | 31 | 34 | 142 | 67 | 48 | 15 | 88 | 27 |
| 4 | 40 | 41 | 3 | 147 | 29 | 56 | 26 | 144 | 7 |
| 5 | 41 | 34 | 13 | 171 | 71 | 70 | 29 | 72 | 56 |
| 6 | 48 | 17 | 17 | 221 | 109 | 52 | 30 | 45 | 29 |
| 7 | 59 | 64 | 28 | 222 | 35 | 30 | 37 | 5 | 183 |
| 8 | 60 | 31 | 83 | 230 | 97 | 31 | 39 | 58 | 17 |
| 9 | 65 | 37 | 65 | 239 | 76 | 13 | 49 | 81 | 34 |
| 10 | 76 | 72 | 21 | 241 | 65 | 37 | 61 | 61 | 37 |
| 11 | 79 | 17 | 39 | 242 | 37 | 25 | 70 | 41 | 41 |
| 12 | 84 | 50 | 59 | 243 | 52 | 32 | 89 | 32 | 51 |
| 13 | 86 | 40 | 40 | 247 | 79 | 51 | 91 | 17 | 16 |
| 14 | 93 | 30 | 40 | 256 | 58 | 38 | 95 | 33 | 29 |
| 15 | 100 | 19 | 17 | 258 | 51 | 55 | 98 | 68 | 20 |
| 16 | 101 | 50 | 36 | 265 | 65 | 64 | 107 | 15 | 52 |
| 17 | 102 | 40 | 34 | 270 | 109 | 33 | 109 | 42 | 18 |
| 18 | 106 | 121 | 18 | 273 | 80 | 31 | 112 | 21 | 81 |
| 19 | 108 | 69 | 18 | 276 | 44 | 134 | 119 | 63 | 37 |
| 20 | 119 | 33 | 24 | 278 | 207 | 0 | 120 | 31 | 56 |
| 21 | 125 | 149 | 13 | 279 | 111 | 20 | 127 | 40 | 82 |
| 22 | 126 | 70 | 39 | 285 | 68 | 19 | 150 | 30 | 47 |
| 23 | 127 | 87 | 19 | 292 | 80 | 35 | 156 | 50 | 52 |
| 24 | 128 | 48 | 65 | 295 | 142 | 11 | 157 | 45 | 46 |
| 25 | 130 | 109 | 35 | 296 | 43 | 34 | 161 | 52 | 34 |
| 26 | 132 | 107 | 64 | 300 | 15 | 83 | 163 | 15 | 50 |
| 27 | 135 | 95 | 32 | 301 | 55 | 39 | 165 | 72 | 42 |
| 28 | 165 | 113 | 14 | 306 | 59 | 56 | 166 | 40 | 55 |
| 29 | 169 | 45 | 67 | 307 | 110 | 16 | 168 | 72 | 42 |
| 30 | 171 | 26 | 13 | 308 | 187 | 0 | 174 | 68 | 48 |
| **Average** | | **61.4** | **34.4** | | **77.2** | **39.2** | | **49.5** | **44.7** |
| **Stdev** | | **36.36** | **20.38** | | **42.93** | **26.60** | | **27.78** | **31.45** |

**Table S2:** Time spent by *N. longicornis* males on hosts with conspecifics and hosts with heterospecifics in the cafeteria assay, having three different categories.

| **S.no.** | **Male ID** | **Time spent on hosts by NL male (sec)** | | **Male ID** | **Time spent on hosts by NL male (sec)** | | **Male ID** | **Time spent on hosts by NL male (sec)** | |
|---|---|---|---|---|---|---|---|---|---|
| | | **Hosts with conspecifics (NL)** | **Hosts with heterospecifics (NV)** | | **Hosts with conspecifics (NL)** | **Hosts with heterospecifics (NG)** | | **Hosts with conspecifics (NL)** | **Hosts with heterospecifics (NO)** |
| 1 | 2 | 62 | 42 | 1 | 73 | 42 | 2 | 58 | 24 |
| 2 | 6 | 72 | 35 | 1.1 | 10 | 171 | 3 | 54 | 57 |
| 3 | 8 | 23 | 126 | 11 | 162 | 9 | 5 | 64 | 54 |
| 4 | 12 | 21 | 114 | 27 | 139 | 18 | 7 | 75 | 43 |
| 5 | 13 | 187 | 28 | 33 | 44 | 71 | 8 | 51 | 41 |
| 6 | 15 | 74 | 30 | 35 | 41 | 10 | 10 | 111 | 30 |
| 7 | 16 | 74 | 88 | 39 | 136 | 7 | 16 | 68 | 55 |
| 8 | 26 | 66 | 68 | 40 | 44 | 71 | 17 | 13 | 131 |
| 9 | 27 | 0 | 186 | 41 | 101 | 39 | 18 | 61 | 75 |
| 10 | 30 | 86 | 22 | 42 | 65 | 65 | 19 | 51 | 51 |
| 11 | 34 | 43 | 47 | 43 | 59 | 69 | 21 | 105 | 45 |
| 12 | 35 | 0 | 38 | 44 | 17 | 88 | 23 | 82 | 17 |
| 13 | 38 | 30 | 67 | 45 | 50 | 69 | 24 | 69 | 48 |
| 14 | 40 | 52 | 82 | 46 | 124 | 6 | 25 | 75 | 31 |
| 15 | 43 | 53 | 73 | 47 | 83 | 40 | 26 | 60 | 59 |
| 16 | 45 | 54 | 77 | 48 | 32 | 122 | 28 | 44 | 86 |
| 17 | 48 | 51 | 24 | 49 | 23 | 12 | 30 | 71 | 55 |
| 18 | 59 | 39 | 123 | 50 | 22 | 107 | 31 | 85 | 14 |
| 19 | 60 | 92 | 17 | 51 | 52 | 97 | 32 | 138 | 25 |
| 20 | 62 | 98 | 71 | 52 | 9 | 56 | 33 | 56 | 57 |
| 21 | 67 | 53 | 117 | 53 | 176 | 10 | 34 | 72 | 43 |
| 22 | 68 | 93 | 67 | 54 | 189 | 0 | 35 | 48 | 40 |
| 23 | 70 | 63 | 57 | 55 | 38 | 70 | 36 | 65 | 58 |
| 24 | 72 | 68 | 72 | 56 | 132 | 8 | 37 | 55 | 49 |
| 25 | 78 | 92 | 43 | 57 | 46 | 60 | 40 | 117 | 28 |
| 26 | 81 | 15 | 49 | 58 | 99 | 52 | 41 | 98 | 4 |
| 27 | 82 | 73 | 59 | 59 | 17 | 103 | 42 | 87 | 24 |
| 28 | 84 | 46 | 52 | 60 | 64 | 45 | 43 | 71 | 38 |
| 29 | 88 | 77 | 54 | 61 | 81 | 57 | 44 | 61 | 85 |
| 30 | 91 | 103 | 53 | 62 | 43 | 91 | 45 | 73 | 39 |
| **Average** | | **62** | **66.03** | | **72.37** | **55.5** | | **71.27** | **46.87** |
| **Stdev** | | **36.41** | **37.19** | | **51.19** | **40.87** | | **24.55** | **24.93** |

**Table S3:** Time spent by *N. giraulti* males on hosts with conspecifics and hosts with heterospecifics in the cafeteria assay, having three different categories.

| S.no. | Male ID | Time spent on hosts by NG male (sec) | | Male ID | Time spent on hosts by NG male (sec) | | Male ID | Time spent on hosts by NG male (sec) | |
|---|---|---|---|---|---|---|---|---|---|
| | | Hosts with conspecifics (NG) | Hosts with heterospecifics (NV) | | Hosts with conspecifics (NG) | Hosts with heterospecifics (NO) | | Hosts with conspecifics (NG) | Hosts with heterospecifics (NL) |
| 1 | 4 | 55 | 56 | 2 | 42 | 85 | 1 | 24 | 34 |
| 2 | 5 | 54 | 35 | 3 | 59 | 101 | 2 | 39 | 132 |
| 3 | 10 | 48 | 52 | 7 | 56 | 48 | 4 | 55 | 56 |
| 4 | 22 | 74 | 49 | 8 | 66 | 79 | 5 | 18 | 137 |
| 5 | 24 | 53 | 90 | 9 | 54 | 61 | 6 | 14 | 43 |
| 6 | 26 | 66 | 64 | 11 | 30 | 61 | 7 | 27 | 47 |
| 7 | 28 | 71 | 71 | 12 | 32 | 31 | 10 | 71 | 37 |
| 8 | 37 | 71 | 50 | 16 | 81 | 42 | 12 | 36 | 66 |
| 9 | 40 | 89 | 42 | 20 | 47 | 51 | 14 | 41 | 64 |
| 10 | 42 | 56 | 51 | 26 | 93 | 37 | 18 | 29 | 35 |
| 11 | 43 | 14 | 141 | 27 | 72 | 56 | 19 | 43 | 66 |
| 12 | 45 | 36 | 71 | 28 | 51 | 90 | 24 | 80 | 43 |
| 13 | 50 | 25 | 58 | 29 | 81 | 56 | 25 | 56 | 56 |
| 14 | 51 | 98 | 39 | 30 | 50 | 60 | 30 | 44 | 39 |
| 15 | 52 | 72 | 40 | 31 | 57 | 60 | 31 | 43 | 31 |
| 16 | 53 | 42 | 35 | 32 | 21 | 156 | 32 | 34 | 51 |
| 17 | 54 | 59 | 55 | 33 | 95 | 34 | 34 | 27 | 44 |
| 18 | 55 | 32 | 28 | 34 | 73 | 67 | 37 | 56 | 41 |
| 19 | 56 | 27 | 40 | 35 | 77 | 38 | 38 | 36 | 68 |
| 20 | 57 | 49 | 76 | 36 | 58 | 43 | 39 | 37 | 86 |
| 21 | 58 | 135 | 13 | 37 | 37 | 37 | 42 | 65 | 58 |
| 22 | 59 | 36 | 93 | 39 | 81 | 32 | 45 | 47 | 56 |
| 23 | 60 | 32 | 83 | 40 | 27 | 53 | 46 | 70 | 62 |
| 24 | 61 | 53 | 48 | 41 | 40 | 79 | 48 | 57 | 77 |
| 25 | 62 | 96 | 41 | 42 | 65 | 63 | 49 | 57 | 45 |
| 26 | 63 | 43 | 58 | 43 | 78 | 53 | 51 | 125 | 25 |
| 27 | 64 | 53 | 50 | 44 | 57 | 44 | 54 | 61 | 33 |
| 28 | 65 | 56 | 41 | 45 | 82 | 42 | 55 | 54 | 66 |
| 29 | 67 | 48 | 67 | 46 | 76 | 59 | 57 | 45 | 67 |
| 30 | 68 | 41 | 67 | 47 | 51 | 51 | 58 | 66 | 57 |
| **Average** | | **56.13** | **56.8** | | **59.63** | **58.97** | | **48.57** | **57.4** |
| **Stdev** | | **25.02** | **24.08** | | **19.84** | **25.35** | | **21.81** | **25.54** |

**Table S4:** Time spent by *N. oneida* males on hosts with conspecifics and hosts with heterospecifics in the cafeteria assay, having three different categories.

| S.no. | Male ID | Time spent on hosts by NO male (sec) | | Male ID | Time spent on hosts by NO male (sec) | | Male ID | Time spent on hosts by NO male (sec) | |
|---|---|---|---|---|---|---|---|---|---|
| | | Hosts with conspecifics (NO) | Hosts with heterospecifics (NV) | | Hosts with conspecifics (NO) | Hosts with heterospecifics (NG) | | Hosts with conspecifics (NO) | Hosts with heterospecifics (NL) |
| 1 | 3 | 23 | 46 | 3 | 73 | 22 | 1 | 47 | 42 |
| 2 | 5 | 17 | 40 | 9 | 78 | 60 | 2 | 69 | 63 |
| 3 | 19 | 38 | 72 | 11 | 51 | 57 | 3 | 56 | 39 |
| 4 | 20 | 59 | 62 | 12 | 55 | 65 | 4 | 44 | 46 |
| 5 | 24 | 66 | 54 | 13 | 38 | 61 | 5 | 25 | 42 |
| 6 | 26 | 43 | 43 | 14 | 18 | 26 | 6 | 40 | 26 |
| 7 | 30 | 60 | 54 | 16 | 59 | 66 | 7 | 48 | 44 |
| 8 | 32 | 3 | 118 | 22 | 56 | 57 | 8 | 57 | 61 |
| 9 | 41 | 58 | 60 | 24 | 48 | 55 | 9 | 20 | 56 |
| 10 | 46 | 82 | 60 | 26 | 42 | 62 | 10 | 38 | 34 |
| 11 | 51 | 36 | 110 | 27 | 76 | 46 | 11 | 38 | 11 |
| 12 | 52 | 37 | 72 | 30 | 19 | 68 | 12 | 50 | 52 |
| 13 | 53 | 44 | 101 | 33 | 52 | 59 | 13 | 116 | 19 |
| 14 | 61 | 70 | 54 | 36 | 80 | 29 | 14 | 54 | 53 |
| 15 | 62 | 50 | 51 | 37 | 55 | 61 | 15 | 140 | 14 |
| 16 | 63 | 81 | 37 | 38 | 56 | 59 | 16 | 41 | 79 |
| 17 | 64 | 34 | 66 | 41 | 30 | 74 | 17 | 55 | 80 |
| 18 | 65 | 25 | 43 | 47 | 24 | 118 | 18 | 75 | 38 |
| 19 | 66 | 34 | 58 | 51 | 56 | 93 | 19 | 32 | 29 |
| 20 | 67 | 56 | 54 | 54 | 94 | 59 | 20 | 37 | 25 |
| 21 | 68 | 46 | 52 | 59 | 9 | 170 | 21 | 17 | 61 |
| 22 | 69 | 60 | 54 | 67 | 63 | 47 | 22 | 42 | 44 |
| 23 | 70 | 8 | 35 | 72 | 42 | 122 | 23 | 71 | 32 |
| 24 | 71 | 84 | 46 | 77 | 80 | 28 | 24 | 42 | 92 |
| 25 | 72 | 36 | 44 | 84 | 60 | 64 | 25 | 55 | 64 |
| 26 | 74 | 45 | 54 | 85 | 50 | 56 | 26 | 42 | 58 |
| 27 | 77 | 46 | 80 | 87 | 51 | 64 | 27 | 36 | 75 |
| 28 | 78 | 45 | 117 | 90 | 58 | 75 | 28 | 57 | 10 |
| 29 | 79 | 99 | 20 | 99 | 33 | 59 | 29 | 70 | 50 |
| 30 | 80 | 18 | 130 | 101 | 23 | 55 | 30 | 41 | 41 |
| **Average** | | **46.77** | **62.9** | | **50.97** | **64.57** | | **51.83** | **46** |
| **Stdev** | | **22.81** | **26.85** | | **20.60** | **29.59** | | **25.24** | **20.79** |

**Python script**: The general structure of the script written in Python used to track males in the cafeteria assay to calculate the distance travelled while searching.

```
#General Structure of Code
import cv2
import numpy as np

input_video = "video.wmv"
with open (file1, "w") as file:
 while cap.isOpened():
   #Code here....
   diff = cv2.absdiff(frame1, frame2)
   gray = cv2.cvtColor(diff, cv2.COLOR_BGR2GRAY)
   blur = cv2.GaussianBlur(gray, (11,11), 0)
   _, thresh = cv2.threshold(blur, 20, 255, cv2.THRESH_BINARY)
   dilated = cv2.dilate(thresh, None, iterations=10)
   contours, _ = cv2.findContours(dilated, cv2.RETR_TREE,  cv2.CHAIN_APPROX_SIMPLE)
   if j>0:
    for contour in contours:
     (x, y, w, h) = cv2.boundingRect(contour)
     if cv2.contourArea(contour) >800:
       continue
     cv2.rectangle(frame1, (x, y), (x+w, y+h), (0, 255, 0), 2)
     cv2.putText(frame1, "Status: {}".format('Movement'), (10, 20),
cv2.FONT_HERSHEY_SIMPLEX,1, (0, 0, 255), 2)
   #Add lines for record track coordinates.
    #..................................
   if cv2.waitKey(15) & 0xFF == ord('q'):
       break
file.close()
cv2.destroyAllWindows()
cap.release()
out.release()
```

**Table S5:** Search distance travelled by *Nasonia* males of all four species in the cafeteria assay looking for the hosts.

| | Search distance (m) | | | |
|---|---|---|---|---|
| S.NO | NV male | NG male | NO male | NL male |
| 1 | 2.43531 | 1.35488 | 1.50544 | 1.08465 |
| 2 | 1.95111 | 0.75846 | 1.9305 | 1.6633 |
| 3 | 1.85135 | 1.32663 | 0.62576 | 0.60637 |
| 4 | 1.55083 | 0.98179 | 0.92735 | 1.00603 |
| 5 | 1.10766 | 0.84583 | 1.23451 | 0.16477 |
| 6 | 1.76983 | 1.47383 | 1.31886 | 0.95305 |
| 7 | 1.56321 | 1.08176 | 1.45151 | 0.28884 |
| 8 | 2.14822 | 1.57797 | 0.29771 | 0.62778 |
| 9 | 1.83271 | 1.89977 | 0.9193 | 0.3941 |
| 10 | 0.74225 | 1.74129 | 0.90983 | 0.8632 |
| 11 | 2.33408 | 1.27617 | 0.80472 | 0.59472 |
| 12 | 2.27015 | 1.68082 | 1.54842 | 0.70401 |
| 13 | 0.91833 | 1.35318 | 0.97843 | 0.68897 |
| 14 | 1.5046 | 1.28386 | 1.70811 | 0.57526 |
| 15 | 0.44789 | 1.23927 | 0.59836 | 0.91072 |
| 16 | 0.25694 | 1.89975 | 1.82854 | 0.62943 |
| 17 | 0.93066 | 1.29117 | 0.6037 | 0.24322 |
| 18 | 2.1396 | 1.3574 | 1.05645 | 0.7957 |
| 19 | 0.93106 | 1.42924 | 1.19833 | 0.7796 |
| 20 | 1.21459 | 1.20192 | 2.33059 | 0.94236 |
| 21 | 2.40709 | 1.62586 | 1.68569 | 0.77306 |
| 22 | 1.3432 | 1.49465 | 1.66139 | 0.69971 |
| 23 | 2.70358 | 1.54559 | 2.17172 | 1.01687 |
| 24 | 1.04187 | 1.32785 | 1.18907 | 0.59119 |
| 25 | 1.26324 | 1.7543 | 2.15142 | 0.47461 |
| 26 | 1.96189 | 1.41088 | 1.04944 | 0.49049 |
| 27 | 1.43484 | 1.70175 | 1.91177 | 0.76993 |
| 28 | 2.0679 | 1.39026 | 1.23141 | 0.92309 |
| 29 | 2.76905 | 1.39336 | 1.30667 | 0.56935 |
| 30 | 2.11695 | 1.40808 | 1.0736 | 1.7714 |
| 31 | 1.83228 | 1.13148 | 1.28925 | 1.15406 |
| 32 | 1.32791 | 1.59888 | 1.34047 | 1.34804 |
| 33 | 1.94703 | 1.20262 | 1.2559 | 1.31228 |
| 34 | 2.32385 | 0.73847 | 1.51152 | 0.61217 |
| 35 | 2.61033 | 0.83414 | 1.90261 | 0.92774 |
| 36 | 2.50641 | 1.43795 | 2.05873 | 0.82543 |
| 37 | 2.03942 | 1.7914 | 1.63456 | 0.57196 |
| 38 | 2.32096 | 1.68414 | 1.11337 | 0.72391 |
| 39 | 1.76601 | 0.67455 | 1.0212 | 0.94892 |
| 40 | 1.45472 | 1.35722 | 0.74243 | 0.26618 |
| 41 | 1.64535 | 1.02994 | 0.3916 | 1.28775 |

Table S5 continued…

| | | | | |
|---|---|---|---|---|
| 42 | 1.25639 | 1.42607 | 1.88839 | 0.66633 |
| 43 | 1.39241 | 0.32502 | 0.58744 | 0.12517 |
| 44 | 0.83858 | 1.72047 | 1.6337 | 0.86171 |
| 45 | 1.4742 | 1.89958 | 1.59647 | 1.28897 |
| 46 | 1.18071 | 1.68338 | 1.63115 | 0.62353 |
| 47 | 2.42931 | 1.82061 | 0.96932 | 0.22321 |
| 48 | 2.65329 | 1.53348 | 1.1166 | 0.79267 |
| 49 | 1.78464 | 0.97249 | 1.34982 | 2.13533 |
| 50 | 1.50954 | 0.45467 | 0.63313 | 2.13128 |
| 51 | 0.52801 | 1.53613 | 1.27823 | 1.07115 |
| 52 | 1.88054 | 0.87445 | 1.51311 | 1.67794 |
| 53 | 0.1191 | 0.36427 | 1.75482 | 1.92965 |
| 54 | 2.58517 | 1.14385 | 0.40994 | 0.92628 |
| 55 | 1.43255 | 0.84518 | 1.34799 | 1.65503 |
| 56 | 2.12339 | 1.42093 | 1.19103 | 1.90898 |
| 57 | 2.42785 | 1.55428 | 0.9485 | 0.97429 |
| 58 | 1.89275 | 0.85205 | 1.68314 | 1.16911 |
| 59 | 1.85139 | 0.57858 | 1.23153 | 1.35472 |
| 60 | 0.74455 | 1.43407 | 0.85147 | 1.38781 |
| 61 | 0.58909 | 0.94607 | 0.69122 | 1.51448 |
| 62 | 0.48543 | 0.56615 | 1.75643 | 0.8802 |
| 63 | 1.34176 | 1.66249 | 2.0725 | 0.86762 |
| 64 | 1.18433 | 1.43665 | 2.14517 | 2.03701 |
| 65 | 1.51078 | 1.44643 | 1.64118 | 1.62552 |
| 66 | 2.63301 | 1.45242 | 1.61361 | 1.0456 |
| 67 | 2.27795 | 1.45376 | 0.88958 | 1.41966 |
| 68 | 2.39705 | 0.87235 | 1.33933 | 1.10407 |
| 69 | 1.26363 | 1.08608 | 1.18611 | 0.53315 |
| 70 | 2.10098 | 1.59533 | 0.8862 | 1.16781 |
| 71 | 1.72443 | 1.16288 | 0.9777 | 1.55193 |
| 72 | 1.85717 | 0.74658 | 1.13298 | 1.2214 |
| **Average** | **1.670198** | **1.284099** | **1.297473** | **0.978414** |
| **Stdev** | **0.647024** | **0.383989** | **0.472091** | **0.48088** |

**Table S6**: Search time used by *Nasonia* males of all four species in the choice assay looking for the hosts.

| | Search time (Sec) | | | |
|---|---|---|---|---|
| S.NO | NV male | NG male | NO male | NL male |
| 1 | 143 | 129 | 171 | 136 |
| 2 | 175 | 151 | 183 | 133 |
| 3 | 196 | 140 | 130 | 91 |
| 4 | 193 | 117 | 119 | 105 |
| 5 | 206 | 97 | 120 | 25 |
| 6 | 148 | 110 | 119 | 136 |
| 7 | 126 | 98 | 122 | 78 |
| 8 | 138 | 119 | 98 | 106 |
| 9 | 147 | 133 | 94 | 54 |
| 10 | 184 | 163 | 139 | 132 |
| 11 | 131 | 126 | 122 | 150 |
| 12 | 160 | 180 | 140 | 106 |
| 13 | 170 | 173 | 172 | 114 |
| 14 | 204 | 115 | 148 | 109 |
| 15 | 166 | 92 | 130 | 165 |
| 16 | 101 | 111 | 142 | 78 |
| 17 | 153 | 125 | 126 | 70 |
| 18 | 131 | 139 | 197 | 80 |
| 19 | 96 | 103 | 110 | 100 |
| 20 | 69 | 139 | 160 | 105 |
| 21 | 113 | 137 | 114 | 176 |
| 22 | 113 | 143 | 78 | 142 |
| 23 | 128 | 125 | 121 | 109 |
| 24 | 201 | 132 | 92 | 84 |
| 25 | 168 | 113 | 145 | 59 |
| 26 | 125 | 80 | 141 | 69 |
| 27 | 155 | 136 | 196 | 83 |
| 28 | 99 | 95 | 115 | 125 |
| 29 | 175 | 149 | 127 | 97 |
| 30 | 151 | 177 | 137 | 125 |
| 31 | 138 | 117 | 136 | 100 |
| 32 | 178 | 142 | 118 | 112 |
| 33 | 156 | 110 | 153 | 135 |
| 34 | 110 | 112 | 129 | 110 |
| 35 | 144 | 99 | 131 | 117 |
| 36 | 134 | 130 | 124 | 86 |
| 37 | 98 | 123 | 125 | 205 |
| 38 | 129 | 63 | 136 | 111 |
| 39 | 62 | 111 | 98 | 91 |
| 40 | 33 | 125 | 91 | 54 |
| 41 | 109 | 139 | 61 | 51 |

Table S6 continued…

| | | | | |
|---|---|---|---|---|
| 42 | 153 | 166 | 130 | 100 |
| 43 | 87 | 160 | 76 | 134 |
| 44 | 163 | 121 | 132 | 89 |
| 45 | 142 | 112 | 116 | 120 |
| 46 | 146 | 109 | 134 | 131 |
| 47 | 125 | 116 | 107 | 102 |
| 48 | 114 | 105 | 162 | 106 |
| 49 | 152 | 182 | 150 | 158 |
| 50 | 125 | 69 | 173 | 129 |
| 51 | 89 | 129 | 174 | 122 |
| 52 | 166 | 85 | 148 | 148 |
| 53 | 52 | 183 | 122 | 117 |
| 54 | 165 | 166 | 191 | 96 |
| 55 | 125 | 132 | 138 | 104 |
| 56 | 157 | 138 | 133 | 138 |
| 57 | 207 | 135 | 86 | 90 |
| 58 | 178 | 131 | 120 | 123 |
| 59 | 152 | 117 | 105 | 134 |
| 60 | 173 | 157 | 127 | 110 |
| 61 | 180 | 166 | 179 | 114 |
| 62 | 138 | 155 | 178 | 141 |
| 63 | 140 | 117 | 162 | 77 |
| 64 | 153 | 137 | 154 | 127 |
| 65 | 118 | 108 | 137 | 152 |
| 66 | 163 | 106 | 106 | 117 |
| 67 | 138 | 138 | 121 | 136 |
| 68 | 149 | 90 | 140 | 95 |
| 69 | 154 | 146 | 129 | 138 |
| 70 | 175 | 120 | 173 | 131 |
| 71 | 126 | 128 | 120 | 94 |
| 72 | 124 | 117 | 158 | 128 |
| **Average** | **141.4583** | **127.2083** | **133.2083** | **111.3194** |
| **Stdev** | **35.61094** | **25.96473** | **28.55878** | **30.59358** |

**Table S7:** Search speed of *Nasonia* males of all four species in the choice assay looking for the hosts.

| | Search speed (m/s) | | | |
|---|---|---|---|---|
| **S.NO** | **NV male** | **NG male** | **NO male** | **NL male** |
| 1 | 0.01174 | 0.00861 | 0.00789 | 0.01351 |
| 2 | 0.01208 | 0.00568 | 0.00346 | 0.01652 |
| 3 | 0.00593 | 0.0113 | 0.00983 | 0.00878 |
| 4 | 0.01133 | 0.0092 | 0.01272 | 0.01134 |
| 5 | 0.00229 | 0.00244 | 0.01462 | 0.01649 |
| 6 | 0.01567 | 0.00646 | 0.00344 | 0.00965 |
| 7 | 0.01146 | 0.00722 | 0.01105 | 0.01591 |
| 8 | 0.01369 | 0.01001 | 0.01215 | 0.01383 |
| 9 | 0.01498 | 0.01413 | 0.01009 | 0.01083 |
| 10 | 0.01352 | 0.00761 | 0.01211 | 0.0095 |
| 11 | 0.01173 | 0.00584 | 0.01009 | 0.01011 |
| 12 | 0.01063 | 0.01103 | 0.00608 | 0.01262 |
| 13 | 0.01218 | 0.00769 | 0.00402 | 0.01328 |
| 14 | 0.0043 | 0.00899 | 0.01187 | 0.00624 |
| 15 | 0.00327 | 0.01498 | 0.01594 | 0.01127 |
| 16 | 0.00352 | 0.01149 | 0.01511 | 0.01604 |
| 17 | 0.00958 | 0.01041 | 0.01303 | 0.01069 |
| 18 | 0.00774 | 0.00875 | 0.00819 | 0.00894 |
| 19 | 0.0128 | 0.00909 | 0.00809 | 0.01044 |
| 20 | 0.01615 | 0.00721 | 0.00837 | 0.01162 |
| 21 | 0.01651 | 0.0097 | 0.0104 | 0.00386 |
| 22 | 0.01609 | 0.01464 | 0.01136 | 0.00891 |
| 23 | 0.00821 | 0.01002 | 0.00808 | 0.01651 |
| 24 | 0.01201 | 0.00711 | 0.01232 | 0.00954 |
| 25 | 0.00883 | 0.0136 | 0.01484 | 0.00804 |
| 26 | 0.01121 | 0.00934 | 0.00744 | 0.00711 |
| 27 | 0.00732 | 0.01216 | 0.00975 | 0.00928 |
| 28 | 0.01071 | 0.01188 | 0.01071 | 0.00738 |
| 29 | 0.01344 | 0.01436 | 0.01029 | 0.00587 |
| 30 | 0.0143 | 0.0128 | 0.00784 | 0.01417 |
| 31 | 0.01454 | 0.01155 | 0.00948 | 0.01154 |
| 32 | 0.00962 | 0.01344 | 0.01136 | 0.01204 |
| 33 | 0.01325 | 0.00904 | 0.00821 | 0.00972 |
| 34 | 0.01263 | 0.00453 | 0.01172 | 0.00557 |
| 35 | 0.01993 | 0.00662 | 0.01452 | 0.00793 |
| 36 | 0.01567 | 0.00799 | 0.0166 | 0.0096 |
| 37 | 0.012 | 0.01035 | 0.01308 | 0.00279 |
| 38 | 0.01138 | 0.01464 | 0.00819 | 0.00652 |
| 39 | 0.01045 | 0.00733 | 0.01042 | 0.01043 |

Table S7 continued…

| | | | | |
|---|---|---|---|---|
| 40 | 0.01064 | 0.01223 | 0.00816 | 0.00493 |
| 41 | 0.0144 | 0.00824 | 0.00642 | 0.02525 |
| 42 | 0.01075 | 0.01026 | 0.01453 | 0.00666 |
| 43 | 0.01898 | 0.00316 | 0.00773 | 9.34E-04 |
| 44 | 0.0132 | 0.01238 | 0.01238 | 0.00968 |
| 45 | 0.00959 | 0.01387 | 0.01376 | 0.01074 |
| 46 | 0.0145 | 0.01177 | 0.01217 | 0.00476 |
| 47 | 0.01215 | 0.01456 | 0.00906 | 0.00219 |
| 48 | 0.01305 | 0.01162 | 0.00689 | 0.00748 |
| 49 | 0.0145 | 0.00744 | 0.01004 | 0.00798 |
| 50 | 0.01561 | 0.01099 | 0.01116 | 0.01251 |
| 51 | 0.01194 | 0.01028 | 0.0036 | 0.00666 |
| 52 | 0.01566 | 0.01155 | 0.00627 | 0.00958 |
| 53 | 0.00633 | 0.00462 | 0.01012 | 0.00659 |
| 54 | 0.01172 | 0.00888 | 0.00691 | 0.00701 |
| 55 | 0.01133 | 0.0082 | 0.01052 | 0.0037 |
| 56 | 0.01207 | 0.01143 | 0.00224 | 0.00592 |
| 57 | 0.01175 | 0.01407 | 0.01069 | 0.0073 |
| 58 | 0.00675 | 0.01329 | 0.00758 | 0.00654 |
| 59 | 0.01621 | 0.01091 | 0.00766 | 0.00396 |
| 60 | 0.01694 | 0.01071 | 0.01219 | 0.00664 |
| 61 | 0.00937 | 0.00815 | 0.00547 | 0.00604 |
| 62 | 0.01166 | 0.00828 | 0.0096 | 0.00528 |
| 63 | 0.00722 | 0.01059 | 0.00369 | 0.00552 |
| 64 | 0.00779 | 0.01387 | 0.01187 | 0.00807 |
| 65 | 0.00854 | 0.01196 | 0.00441 | 0.00347 |
| 66 | 0.01398 | 0.01281 | 0.00997 | 0.00995 |
| 67 | 0.0107 | 0.01036 | 0.0099 | 0.0078 |
| 68 | 0.00745 | 0.01335 | 0.01665 | 0.00897 |
| 69 | 0.01695 | 0.01114 | 0.01307 | 0.00439 |
| 70 | 0.0092 | 0.01246 | 0.0096 | 0.00493 |
| 71 | 0.02163 | 0.01207 | 0.0181 | 0.00933 |
| 72 | 0.00914 | 0.01135 | 0.00753 | 0.00704 |
| **Average** | **0.011728** | **0.010223** | **0.009926** | **0.00892** |
| **Stdev** | **0.003787** | **0.002868** | **0.003431** | **0.004044** |

**Table S8:** Latency to find the first host with conspecifics (Search latency) by *Nasonia* males of all four species in the choice assay looking for the hosts.

| | Latency to find first host with conspecifics (Search latency) | | | |
|---|---|---|---|---|
| **S.NO** | **NV male** | **NG male** | **NO male** | **NL male** |
| 1 | 4 | 19 | 39 | 46 |
| 2 | 8 | 14 | 35 | 27 |
| 3 | 49 | 25 | 26 | 38 |
| 4 | 23 | 25 | 20 | 16 |
| 5 | 20 | 10 | 9 | 53 |
| 6 | 16 | 12 | 20 | 16 |
| 7 | 53 | 17 | 5 | 12 |
| 8 | 6 | 4 | 13 | 26 |
| 9 | 44 | 4 | 4 | 240 |
| 10 | 40 | 12 | 14 | 57 |
| 11 | 13 | 72 | 19 | 49 |
| 12 | 34 | 24 | 16 | 8 |
| 13 | 62 | 23 | 66 | 74 |
| 14 | 59 | 6 | 62 | 17 |
| 15 | 22 | 49 | 21 | 21 |
| 16 | 27 | 24 | 20 | 15 |
| 17 | 37 | 17 | 17 | 44 |
| 18 | 35 | 24 | 228 | 7 |
| 19 | 13 | 18 | 42 | 48 |
| 20 | 25 | 6 | 83 | 22 |
| 21 | 11 | 17 | 10 | 44 |
| 22 | 18 | 10 | 17 | 24 |
| 23 | 17 | 5 | 21 | 8 |
| 24 | 17 | 24 | 74 | 34 |
| 25 | 10 | 21 | 18 | 15 |
| 26 | 17 | 41 | 4 | 27 |
| 27 | 8 | 18 | 108 | 22 |
| 28 | 7 | 17 | 40 | 41 |
| 29 | 9 | 13 | 50 | 10 |
| 30 | 27 | 7 | 4 | 31 |
| 31 | 3 | 24 | 46 | 41 |
| 32 | 30 | 14 | 18 | 41 |
| 33 | 17 | 6 | 49 | 26 |
| 34 | 97 | 15 | 28 | 31 |
| 35 | 5 | 68 | 28 | 26 |
| 36 | 7 | 13 | 9 | 65 |
| 37 | 47 | 7 | 24 | 91 |
| 38 | 5 | 44 | 36 | 55 |
| 39 | 14 | 4 | 6 | 7 |
| 40 | 23 | 14 | 18 | 49 |

Table S8 continued…

| | | | | |
|---|---|---|---|---|
| 41 | 3 | 6 | 48 | 22 |
| 42 | 29 | 5 | 16 | 10 |
| 43 | 29 | 58 | 28 | 33 |
| 44 | 30 | 48 | 13 | 15 |
| 45 | 67 | 15 | 10 | 210 |
| 46 | 6 | 19 | 33 | 60 |
| 47 | 5 | 5 | 27 | 159 |
| 48 | 13 | 4 | 67 | 40 |
| 49 | 15 | 40 | 46 | 37 |
| 50 | 8 | 6 | 86 | 31 |
| 51 | 30 | 19 | 134 | 20 |
| 52 | 7 | 25 | 60 | 16 |
| 53 | 6 | 19 | 19 | 20 |
| 54 | 21 | 19 | 59 | 32 |
| 55 | 38 | 25 | 49 | 33 |
| 56 | 17 | 16 | 45 | 54 |
| 57 | 51 | 20 | 15 | 57 |
| 58 | 8 | 21 | 74 | 4 |
| 59 | 20 | 9 | 49 | 14 |
| 60 | 225 | 13 | 21 | 31 |
| 61 | 29 | 13 | 16 | 28 |
| 62 | 54 | 18 | 33 | 8 |
| 63 | 24 | 47 | 8 | 11 |
| 64 | 64 | 5 | 26 | 58 |
| 65 | 5 | 24 | 106 | 41 |
| 66 | 3 | 4 | 29 | 39 |
| 67 | 17 | 11 | 6 | 11 |
| 68 | 31 | 11 | 32 | 4 |
| 69 | 21 | 34 | 51 | 20 |
| 70 | 46 | 19 | 84 | 27 |
| 71 | 28 | 12 | 16 | 30 |
| 72 | 38 | 10 | 39 | 35 |
| **Average** | **27.31944** | **19.26389** | **37.66667** | **37.97222** |
| **Stdev** | **30.13623** | **14.67199** | **35.35414** | **39.52926** |

**Table S9:** List of Identified peaks in CHC profile of hosts with adult males and females inside: In total 46 CHC compounds have been found in the extract collected from the parasitized hosts having adult males and females inside. We have included only compounds which were present in all three replicates. The values represent the percentage relative abundance ± standard deviation (S.D.).

| S.No. | Retention time | Diagnostic ions | Compound name | Linear retention index | Mean relative abundance (%) ± S.D. Hosts with wasps | | | |
|---|---|---|---|---|---|---|---|---|
| | | | | | NV | NG | NO | NL |
| 1 | 56.324 | 352 | C25 | 2500 | 0.36±0.06 | 0.4±0.17 | 0.57±0.02 | 0.21±0.04 |
| 2 | 57.694 | 337 | MeC25(3-) | 2573 | 0±0 | 0.12±0.02 | 0.35±0.23 | 0±0 |
| 3 | 58.202 | 366 | C26 | 2600 | 0.23±0.05 | 0.26±0.31 | 0.79±0.35 | 0±0 |
| 4 | 60.029 | 380 | C27 | 2700 | 0.76±0.05 | 0.82±0.25 | 1.6±0.11 | 0.58±0.2 |
| 5 | 60.62 | 22,41,96,168 | MeC27(15-;13-;11-;9-) | 2735 | 0.63±0.06 | 0.74±0.48 | 1.57±0.43 | 0.39±0.15 |
| 6 | 61.331 | 365 | MeC27(3-) | 2773 | 0.47±0.11 | 0.77±0.2 | 1.48±0.04 | 0.38±0.05 |
| 7 | 61.793 | 394 | C28 | 2800 | 0.2±0.05 | 0±0 | 0±0 | 0±0 |
| 8 | 63.5 | 408 | C29 | 2900 | 4.2±0.93 | 2.1±0.5 | 2.03±0.68 | 2.63±0.27 |
| 9 | 64.029 | 22,41,96,168 | MeC29(15-;13-;11-) | 2932 | 1.3±0.26 | 1.49±0.15 | 1.72±0.26 | 0.8±0.23 |
| 10 | 64.18 | 112 | MeC29(7-) | 2941 | 0.94±0.12 | 0.42±0.03 | 0.44±0.09 | 0.32±0.1 |
| 11 | 64.335 | 85 | MeC29(5-) | 2951 | 0.89±0.16 | 0.73±0.04 | 0.58±0.15 | 0.47±0.15 |
| 12 | 64.719 | 393 | MeC29(3-) | 2973 | 0±0 | 2.4±0.53 | 3.64±1.06 | 2.5±0.22 |
| 13 | 64.844 | 196, 85 | DiMeC29(5-, x-) | 2987 | 2.45±0.37 | 0.11±0.05 | 0.38±0.1 | 0.1±0.02 |
| 14 | 65.145 | 422 | C30 | 3000 | 0.93±0.15 | 0.71±0.09 | 0.81±0.04 | 0.33±0.02 |
| 15 | 65.648 | 407,238, 196, 182,154 | DiMeC29(3,11-;3,13-;3,15-) | 3012 | 0.17±0.08 | 0±0 | 0±0 | 0.12±0.03 |
| 16 | 65.798 | 112 | MeC30(7-; 8-) | 3041 | 0.21±0.07 | 0±0 | 0±0 | 0±0 |
| 17 | 66.4 | 55,434 | C31-ene-1 | - | 0.09±0.02 | 0±0 | 0±0 | 0±0 |
| 18 | 66.54 | 55, 434 | C31-ene-2 | - | 0.14±0.03 | 0±0 | 0±0 | 0.1±0.02 |
| 19 | 66.81 | 436 | C31 | 3100 | 18.93±1.08 | 14.29±1.02 | 15.11±0.59 | 12.1±0.22 |
| 20 | 67.236 | 224, 196, 168 | MeC31(15-;13-;11-) | 3131 | 5.75±0.99 | 4.25±0.22 | 3.21±0.2 | 2.46±2.06 |
| 21 | 67.417 | 112, 365 | MeC31(7-) | 3141 | 7.08±0.62 | 3.29±0.24 | 3.01±0.36 | 5.67±0.68 |
| 22 | 67.568 | 85, 393 | MeC31(5-) | 3150 | 3.36±0.42 | 3.89±0.13 | 3.41±0.34 | 2.11±0.24 |
| 23 | 67.838 | 112, 308 | DiMeC31(7,11-) | 3168 | 0.39±0.12 | 0.92±0.22 | 0.75±0.03 | 0.51±0.06 |
| 24 | 67.936 | 421 | MeC31(3-) | 3174 | 3.79±0.45 | 6.95±0.75 | 6.05±0.96 | 9.14±0.36 |
| 25 | 68.18 | 112 | DiMeC31(7, x-) | 3181 | 3.9±0.94 | 1.03±0.12 | 1.83±0.42 | 0.24±0.06 |
| 26 | 68.351 | 450 | C32 | 3200 | 0.75±0.09 | 1.12±0.15 | 0.81±0.07 | 0.43±0.04 |
| 27 | 68.424 | 97 | MeC32(6-) | 3245 | 0.64±0.09 | 0.57±0.24 | 0.21±0.24 | 0.23±0.02 |
| 28 | 68.969 | 393 | TetraMeC31(3,7,11,15-) | 3258 | 0.49±0.09 | 1.26±0.41 | 0±0 | 0.27±0.05 |
| 29 | 69.498 | 55 | C33-ene_1 | - | 0.08±0.01 | 0±0 | 0±0 | 0.4±0.14 |
| 30 | 69.597 | 55 | C33-ene_2 | - | 0.1±0.03 | 0±0 | 0±0 | 1.36±0.2 |
| 31 | 69.794 | 464 | C33 | 3300 | 4.54±0.49 | 2.78±0.41 | 1.67±0.28 | 0±0 |
| 32 | 70.282 | 196, 224, 168 | MeC33(15-;13-;11-) | 3335 | 12.66±1.26 | 17.36±0.92 | 17.92±0.47 | 15.66±0.78 |
| 33 | 70.422 | 112, 393 | MeC33(7-) | 3340 | 0.79±0.18 | 0.64±0.13 | 1.43±0.24 | 1.12±0.63 |
| 34 | 70.51 | 85,420 | MeC33(5-) | 3350 | 0.15±0.08 | 2.75±1.36 | 6.95±0.82 | 11.63±0.11 |
| 35 | 70.89 | 239, 168 | DiMeC33(11,15-;11,21-) | 3355 | 1.34±0.33 | 1.41±0.95 | 0.34±0.1 | 0.65±0.14 |

Table S9 continued…

| | | | | | | | | |
|---|---|---|---|---|---|---|---|---|
| 36 | 70.952 | 112 | DiMeC33(7,19-;7,23) | 3368 | 0.48±0.23 | 0±0 | 0±0 | 1.95±0.47 |
| 37 | 71.039 | 449 | MeC33(3-) | 3373 | 0.92±0.26 | 4.19±0.28 | 5.6±0.87 | 0±0 |
| 38 | 71.407 | 477, 463, 308, 280, 266, 238 | DiMeC33(3,17-;3,15-) | 3395 | 1.5±0.09 | 2.21±0.13 | 1.96±0.27 | 1.29±0.31 |
| 39 | 71.828 | 478 | C34 | 3400 | 0.45±0.08 | 0±0 | 0±0 | 0.67±0.06 |
| 40 | 71.957 | 491, 421, 351, 266, 197, 127 | TetraMeC33(3,7,11,15-) | 3460 | 0.12±0.05 | 6.36±1.09 | 1.82±0.31 | 0.33±0.04 |
| 41 | 72.38 | 492 | C35 | 3500 | 1.37±0.42 | 0±0 | 0±0 | 2.33±0.44 |
| 42 | 73.706 | 252, 491 | MeC35(17-) | 3530 | 0±0 | 8.64±3.92 | 3.57±0.23 | 4.22±0.56 |
| 43 | 74.25 | 224, 196, 168 | MeC35(15-;13-;11-) | 3532 | 5.95±0.33 | 4.03±2.43 | 6.92±0.63 | 14.07±0.89 |
| 44 | 74.531 | 112 | MeC35(7-) | 3542 | 8.12±0.61 | 0±0 | 0±0 | 0.29±0.12 |
| 45 | 74.754 | 477, 463, 85 | MeC35(5-;3-) | 3560 | 1.26±0.52 | 0.99±1.05 | 1.49±0.28 | 1.98±0.62 |
| 46 | 78.543 | 196, 224 | MeC37(15-;13-) | 3730 | 1.11±0.15 | 0±0 | 0±0 | 0±0 |

**Table S10**: List of Identified peaks in CHC profile of adult females: In total 50 compounds have been found in the extract collected from the adult females. We have included only compounds which were present in all three replicates. The values represent the percentage relative abundance ± standard deviation (S.D.).

| S.NO. | Retention time | Diagnostic ions | Compound name | Linear retention index | Mean relative abundance (%) ± S.D. | | | |
|---|---|---|---|---|---|---|---|---|
| | | | | | NV Female | NG Female | NO Female | NL Female |
| 1 | 56.324 | 352 | C25 | 2500 | 0.27±0.09 | 0.27±0.28 | 0±0 | 0.27±0.35 |
| 2 | 57.294 | 85 | MeC25(5-) | 2551 | 0.2±0.04 | 0.07±0.04 | 0±0 | 0.06±0.01 |
| 3 | 57.694 | 337 | MeC25(3-) | 2573 | 0.03±0.01 | 0.14±0.19 | 0±0 | 0.06±0 |
| 4 | 58.202 | 366 | C26 | 2600 | 0.01±0 | 0.02±0.01 | 0±0 | 0.13±0.01 |
| 5 | 60.029 | 380 | C27 | 2700 | 1.16±0.34 | 0.85±0.82 | 0±0 | 0.53±0.03 |
| 6 | 60.62 | 22,41,96,168 | MeC27(15-;13-;11-;9-) | 2735 | 0.02±0.01 | 0.06±0.06 | 0±0 | 0.05±0.01 |
| 7 | 60.76 | 112 | MeC27(7-) | 2740 | 0.04±0.02 | 0.02±0.01 | 0±0 | 0.02±0 |
| 8 | 60.931 | 85 | MeC27(5-) | 2751 | 0.29±0.08 | 0.08±0.05 | 0±0 | 0.3±0 |
| 9 | 61.331 | 365 | MeC27(3-) | 2773 | 0.13±0.07 | 0.28±0.18 | 0±0 | 0.08±0 |
| 10 | 61.793 | 394 | C28 | 2800 | 0.13±0.03 | 0.07±0.07 | 0±0 | 0.15±0 |
| 11 | 62.794 | 365 | MeC28(4-) | 2858 | 0±0 | 0±0 | 0±0 | 0.02±0 |
| 12 | 63.5 | 408 | C29 | 2900 | 12.24±0.62 | 2.66±0.81 | 3±0.07 | 11.31±0.16 |
| 13 | 64.029 | 22,41,96,168 | MeC29(15-;13-;11-) | 2932 | 0.74±0.1 | 0.42±0.12 | 0.13±0.06 | 0.96±0.1 |
| 14 | 64.18 | 112 | MeC29(7-) | 2941 | 2.66±0.33 | 0.28±0.08 | 0.19±0.06 | 1.24±0.05 |
| 15 | 64.335 | 85 | MeC29(5-) | 2951 | 1.46±0.19 | 0.54±0.14 | 0.24±0.09 | 0.4±0.51 |
| 16 | 64.719 | 393 | MeC29(3-) | 2973 | 1.7±0.63 | 0.74±0.24 | 0.16±0.13 | 2.52±0.03 |
| 17 | 64.844 | 196, 85 | DiMeC29(5-, x-) | 2987 | 0.15±0.01 | 0.1±0.02 | 0.07±0.01 | 0.12±0.01 |
| 18 | 65.145 | 422 | C30 | 3000 | 1.2±0.19 | 0.85±0.07 | 1.23±0.43 | 1.58±0.09 |
| 19 | 65.648 | 407,238, 196, 182,154 | DiMeC29(3,11-;3,13-;3,15-) | 3012 | 0.12±0.04 | 0.1±0.01 | 0±0 | 0±0 |
| 20 | 65.798 | 112 | MeC30(7-; 8-) | 3041 | 0.25±0.12 | 0.06±0.02 | 0±0 | 0.63±0.06 |
| 21 | 66.4 | 55,434 | C31-ene-1 | - | 0.29±0.13 | 0.06±0.02 | 0±0 | 0.21±0.01 |
| 22 | 66.54 | 55, 434 | C31-ene-2 | - | 0.11±0.04 | 0.04±0.01 | 0±0 | 0.02±0 |
| 23 | 66.81 | 436 | C31 | 3100 | 21.2±1.24 | 15.6±0.48 | 29.83±1.17 | 21.18±1.04 |
| 24 | 67.236 | 224, 196, 168 | MeC31(15-;13-;11-) | 3131 | 6.26±0.38 | 3.53±0.94 | 3.85±0.47 | 2.9±0.12 |
| 25 | 67.417 | 112, 365 | MeC31(7-) | 3141 | 9.24±0.5 | 4.73±0.67 | 4.2±0.46 | 9.05±0.51 |
| 26 | 67.568 | 85, 393 | MeC31(5-) | 3150 | 5.51±0.08 | 5.46±0.42 | 5.79±0.27 | 3.41±0.08 |
| 27 | 67.838 | 112, 308 | DiMeC31(7,11-) | 3168 | 0.41±0.04 | 0.32±0.11 | 5±1.16 | 2.02±0.05 |
| 28 | 67.936 | 421 | MeC31(3-) | 3174 | 2.61±0.33 | 4.69±0.74 | 1.47±1.79 | 1.81±0.04 |
| 29 | 68.18 | 112 | DiMeC31(7, x-) | 3181 | 0.46±0.11 | 0.09±0.01 | 3.44±0.4 | 0.17±0 |
| 30 | 68.351 | 450 | C32 | 3200 | 0.41±0.05 | 1.4±0.07 | 2.02±1.51 | 1.23±0.08 |
| 31 | 68.424 | 97 | MeC32(6-) | 3245 | 0.99±0.13 | 0.22±0.04 | 0.64±0.03 | 0.12±0.01 |
| 32 | 68.969 | 393 | TetraMeC31(3,7,11,15-) | 3258 | 0.16±0.01 | 0.06±0 | 0±0 | 3.68±0.06 |
| 33 | 69.498 | 55 | C33-ene_1 | - | 0.46±0.57 | 0.1±0.04 | 0±0 | 0.24±0.29 |
| 34 | 69.597 | 55 | C33-ene_2 | - | 0.21±0.03 | 0.4±0.1 | 0±0 | 0.1±0.01 |

Table S10 continued…

| | | | | | | | | |
|---|---|---|---|---|---|---|---|---|
| 35 | 69.794 | 464 | C33 | 3300 | 1.05±0.06 | 1.19±0.22 | 0.78±0.11 | 0±0 |
| 36 | 70.282 | 196, 224, 168 | MeC33(15-;13-;11-) | 3335 | 12±0.38 | 1.82±1.2 | 13.95±1.15 | 14.82±0.83 |
| 37 | 70.422 | 112, 393 | MeC33(7-) | 3340 | 0.81±0 | 0.66±0.19 | 1.44±0.97 | 0.38±0.08 |
| 38 | 70.51 | 85,420 | MeC33(5-) | 3350 | 0.72±0.05 | 0±0 | 5.7±0.63 | 0.42±0.03 |
| 39 | 70.89 | 239, 168 | DiMeC33(11,15-;11,21-) | 3355 | 1.66±0.06 | 6.16±1.86 | 0±0 | 0.38±0.49 |
| 40 | 70.952 | 112 | DiMeC33(7,19-;7,23-) | 3368 | 0.25±0.08 | 0.5±0.17 | 0.31±0.03 | 0.73±0.54 |
| 41 | 71.039 | 449 | MeC33(3-) | 3373 | 1.48±0.13 | 8.13±0.83 | 8.43±0.18 | 2.7±0.17 |
| 42 | 71.407 | 477, 463, 308, 280, 266, 238 | DiMeC33(3,17-;3,15-) | 3395 | 3.15±0.19 | 3.52±0.5 | 1.35±0.14 | 0.93±0.07 |
| 43 | 71.828 | 478 | C34 | 3400 | 0.35±0.01 | 0.5±0.08 | 0.21±0.02 | 1.56±0.01 |
| 44 | 71.957 | 491, 421, 351, 266, 197, 127 | TetraMeC33(3,7,11,15-) | 3460 | 0.33±0.11 | 0.63±0.06 | 0.18±0.04 | 0±0 |
| 45 | 72.38 | 492 | C35 | 3500 | 1.66±0.33 | 8.3±0.52 | 1.41±0.4 | 1.54±0.07 |
| 46 | 73.706 | 252, 491 | MeC35(17-) | 3530 | 1.53±0.12 | 4.13±0.86 | 1±0.29 | 2.01±0.04 |
| 47 | 74.25 | 224, 196, 168 | MeC35(15-;13-;11-) | 3532 | 2.64±0.63 | 8.21±0.72 | 3.36±1.15 | 7.07±1.17 |
| 48 | 74.531 | 112 | MeC35(7-) | 3542 | 0.5±0.11 | 0.35±0.1 | 0±0 | 0±0 |
| 49 | 74.754 | 477, 463, 85 | MeC35(5-;3-) | 3560 | 0.73±0.03 | 3.31±0.12 | 0.61±0.28 | 0.91±0.08 |
| 50 | 78.543 | 196, 224 | MeC37(15-;13-) | 3730 | 0±0 | 1.02±0.4 | 0±0 | 0±0 |

**Table S11:** Results of SIMPER analysis of the cuticular hydrocarbon profiles of hosts parasitised by *N. vitripennis* and *N. giraulti* females. Compounds in bold represent compounds that are part of peaks responsible for 50% dissimilarity in females.

| S.No. | Compound name | Average dissimilarity | Contribution % | Cumulative % | Mean Hosts with wasps (NV) | Mean Hosts with wasps (NG) |
|---|---|---|---|---|---|---|
| 1 | MeC35(17-) | 4.321 | 11.94 | 11.94 | 0 | 8.64 |
| 2 | MeC35(7-) | 4.061 | 11.22 | 23.16 | 8.12 | 0 |
| 3 | TetraMeC33(3,7,11,15-) | 3.119 | 8.616 | 31.77 | 0.124 | 6.36 |
| 4 | **MeC33(15-;13-;11-)** | 2.347 | 6.484 | 38.26 | 12.7 | 17.4 |
| 5 | **C31** | 2.321 | 6.411 | 44.67 | 18.9 | 14.3 |
| 6 | MeC31(7-) | 1.896 | 5.237 | 49.9 | 7.08 | 3.29 |
| 7 | **MeC33(3-)** | 1.632 | 4.509 | 54.41 | 0.924 | 4.19 |
| 8 | MeC31(3-) | 1.578 | 4.36 | 58.77 | 3.79 | 6.95 |
| 9 | DiMeC31(7,x-) | 1.435 | 3.963 | 62.74 | 3.9 | 1.03 |
| 10 | MeC33(5-) | 1.3 | 3.592 | 66.33 | 0.147 | 2.75 |
| 11 | MeC35(15-;13-;11-) | 1.246 | 3.441 | 69.77 | 5.95 | 4.03 |
| 12 | MeC29(3-) | 1.201 | 3.319 | 73.09 | 0 | 2.4 |
| 13 | DiMeC29(5-,x-) | 1.167 | 3.223 | 76.31 | 2.45 | 0.114 |
| 14 | C29 | 1.049 | 2.897 | 79.21 | 4.2 | 2.1 |
| 15 | C33 | 0.8775 | 2.424 | 81.63 | 4.54 | 2.78 |
| 16 | MeC31(15-;13-;11-) | 0.7535 | 2.082 | 83.71 | 5.75 | 4.25 |
| 17 | C35 | 0.6844 | 1.891 | 85.6 | 1.37 | 0 |
| 18 | MeC37(15-;13-) | 0.5527 | 1.527 | 87.13 | 1.11 | 0 |
| 19 | MeC35(5-;3-) | 0.4377 | 1.209 | 88.34 | 1.26 | 0.985 |
| 20 | TetraMeC31(3,7,11,15-) | 0.3855 | 1.065 | 89.41 | 0.486 | 1.26 |
| 21 | DiMeC33(11,15-;11,21-) | 0.3769 | 1.041 | 90.45 | 1.34 | 1.41 |
| 22 | DiMeC33(3,17-;3,15-) | 0.3541 | 0.9781 | 91.42 | 1.5 | 2.21 |
| 23 | MeC31(5-) | 0.2716 | 0.7503 | 92.18 | 3.36 | 3.89 |
| 24 | DiMeC31(7,11-) | 0.2631 | 0.7268 | 92.9 | 0.391 | 0.917 |
| 25 | MeC29(7-) | 0.2614 | 0.7221 | 93.62 | 0.943 | 0.42 |
| 26 | DiMeC33(7,19-;7,23-) | 0.2402 | 0.6636 | 94.29 | 0.48 | 0 |
| 27 | C34 | 0.2275 | 0.6284 | 94.92 | 0.455 | 0 |
| 28 | C32 | 0.1865 | 0.5153 | 95.43 | 0.746 | 1.12 |
| 29 | MeC27(15-;13-;11-;9-) | 0.1671 | 0.4617 | 95.89 | 0.634 | 0.742 |
| 30 | MeC27(3-) | 0.1507 | 0.4164 | 96.31 | 0.468 | 0.769 |
| 31 | MeC29(15-;13-;11-) | 0.1251 | 0.3456 | 96.66 | 1.3 | 1.49 |
| 32 | C26 | 0.1153 | 0.3186 | 96.97 | 0.232 | 0.259 |
| 33 | C30 | 0.1107 | 0.3058 | 97.28 | 0.933 | 0.711 |
| 34 | MeC30(7-;8-) | 0.1032 | 0.285 | 97.56 | 0.206 | 0 |
| 35 | C28 | 0.09796 | 0.2706 | 97.84 | 0.196 | 0 |
| 36 | C27 | 0.09617 | 0.2657 | 98.1 | 0.755 | 0.822 |
| 37 | MeC33(7-) | 0.0951 | 0.2627 | 98.36 | 0.787 | 0.638 |
| 38 | MeC32(6-) | 0.09473 | 0.2617 | 98.63 | 0.636 | 0.565 |
| 39 | MeC29(5-) | 0.08813 | 0.2435 | 98.87 | 0.89 | 0.732 |

Table S11 continued…

| | | | | | | |
|---|---|---|---|---|---|---|
| 40 | DiMeC29(3,11-;3,13-;3,15-) | 0.08554 | 0.2363 | 99.1 | 0.171 | 0 |
| 41 | C31-ene-2 | 0.06961 | 0.1923 | 99.3 | 0.139 | 0 |
| 42 | C25 | 0.0631 | 0.1743 | 99.47 | 0.361 | 0.398 |
| 43 | MeC25(3-) | 0.0579 | 0.1599 | 99.63 | 0 | 0.116 |
| 44 | C33-ene_2 | 0.0512 | 0.1414 | 99.77 | 0.102 | 0 |
| 45 | C31-ene-1 | 0.04265 | 0.1178 | 99.89 | 0.0853 | 0 |
| 46 | C33-ene_1 | 0.03957 | 0.1093 | 100 | 0.0791 | 0 |

**Table S12:** Results of SIMPER analysis of the cuticular hydrocarbon profiles of hosts parasitised by *N. vitripennis* and *N. oneida* females. Compounds in bold represent compounds that are part of peaks responsible for 50% dissimilarity in females.

| S.No. | Compound name | Average dissimilarity | Contribution % | Cumulative % | Mean Hosts with wasps (NV) | Mean Hosts with wasps (NO) |
|---|---|---|---|---|---|---|
| 1 | MeC35(7-) | 4.061 | 11.52 | 11.52 | 8.12 | 0 |
| 2 | **MeC33(5-)** | 3.401 | 9.648 | 21.17 | 0.147 | 6.95 |
| 3 | MeC33(15-;13-;11-) | 2.626 | 7.45 | 28.62 | 12.7 | 17.9 |
| 4 | **MeC33(3-)** | 2.337 | 6.629 | 35.25 | 0.924 | 5.6 |
| 5 | **MeC31(7-)** | 2.038 | 5.781 | 41.03 | 7.08 | 3.01 |
| 6 | **C31** | 1.91 | 5.418 | 46.45 | 18.9 | 15.1 |
| 7 | MeC29(3-) | 1.819 | 5.159 | 51.6 | 0 | 3.64 |
| 8 | MeC35(17-) | 1.783 | 5.058 | 56.66 | 0 | 3.57 |
| 9 | C33 | 1.431 | 4.06 | 60.72 | 4.54 | 1.67 |
| 10 | MeC31(15-;13-;11-) | 1.273 | 3.61 | 64.33 | 5.75 | 3.21 |
| 11 | MeC31(3-) | 1.127 | 3.196 | 67.53 | 3.79 | 6.05 |
| 12 | C29 | 1.088 | 3.085 | 70.61 | 4.2 | 2.03 |
| 13 | DiMeC31(7,x-) | 1.037 | 2.941 | 73.56 | 3.9 | 1.83 |
| 14 | DiMeC29(5-,x-) | 1.032 | 2.927 | 76.48 | 2.45 | 0.383 |
| 15 | TetraMeC33(3,7,11,15-) | 0.8491 | 2.409 | 78.89 | 0.124 | 1.82 |
| 16 | C35 | 0.6844 | 1.941 | 80.83 | 1.37 | 0 |
| 17 | MeC37(15-;13-) | 0.5527 | 1.568 | 82.4 | 1.11 | 0 |
| 18 | MeC27(3-) | 0.5057 | 1.435 | 83.83 | 0.468 | 1.48 |
| 19 | DiMeC33(11,15-;11,21-) | 0.5005 | 1.42 | 85.25 | 1.34 | 0.337 |
| 20 | MeC35(15-;13-;11-) | 0.485 | 1.376 | 86.63 | 5.95 | 6.92 |
| 21 | MeC27(15-;13-;11-;9-) | 0.4677 | 1.327 | 87.96 | 0.634 | 1.57 |
| 22 | C27 | 0.4208 | 1.194 | 89.15 | 0.755 | 1.6 |
| 23 | MeC33(7-) | 0.3199 | 0.9075 | 90.06 | 0.787 | 1.43 |
| 24 | C26 | 0.2765 | 0.7842 | 90.84 | 0.232 | 0.785 |
| 25 | MeC29(7-) | 0.2531 | 0.718 | 91.56 | 0.943 | 0.437 |
| 26 | TetraMeC31(3,7,11,15-) | 0.2432 | 0.6899 | 92.25 | 0.486 | 0 |
| 27 | DiMeC33(7,19-;7,23-) | 0.2402 | 0.6814 | 92.93 | 0.48 | 0 |
| 28 | DiMeC33(3,17-;3,15-) | 0.2308 | 0.6547 | 93.59 | 1.5 | 1.96 |
| 29 | C34 | 0.2275 | 0.6453 | 94.23 | 0.455 | 0 |
| 30 | MeC29(15-;13-;11-) | 0.2118 | 0.6008 | 94.83 | 1.3 | 1.72 |
| 31 | MeC32(6-) | 0.2115 | 0.6 | 95.43 | 0.636 | 0.213 |
| 32 | MeC35(5-;3-) | 0.2105 | 0.5971 | 96.03 | 1.26 | 1.49 |
| 33 | MeC31(5-) | 0.1818 | 0.5156 | 96.54 | 3.36 | 3.41 |
| 34 | DiMeC31(7,11-) | 0.1798 | 0.5099 | 97.05 | 0.391 | 0.75 |
| 35 | MeC25(3-) | 0.1735 | 0.4921 | 97.55 | 0 | 0.347 |
| 36 | MeC29(5-) | 0.1583 | 0.4491 | 98 | 0.89 | 0.576 |
| 37 | C25 | 0.107 | 0.3036 | 98.3 | 0.361 | 0.575 |

Table S12 continued…

| | | | | | | |
|---|---|---|---|---|---|---|
| 38 | MeC30(7-;8-) | 0.1032 | 0.2927 | 98.59 | 0.206 | 0 |
| 39 | C28 | 0.09796 | 0.2779 | 98.87 | 0.196 | 0 |
| 40 | DiMeC29(3,11-;3,13-;3,15-) | 0.08554 | 0.2426 | 99.11 | 0.171 | 0 |
| 41 | C31-ene-2 | 0.06961 | 0.1974 | 99.31 | 0.139 | 0 |
| 42 | C30 | 0.0631 | 0.179 | 99.49 | 0.933 | 0.813 |
| 43 | C33-ene_2 | 0.0512 | 0.1452 | 99.63 | 0.102 | 0 |
| 44 | C32 | 0.04663 | 0.1323 | 99.77 | 0.746 | 0.812 |
| 45 | C31-ene-1 | 0.04265 | 0.121 | 99.89 | 0.0853 | 0 |
| 46 | C33-ene_1 | 0.03957 | 0.1122 | 100 | 0.0791 | 0 |

**Table S13:** Results of SIMPER analysis of the cuticular hydrocarbon profiles of hosts parasitised by *N. vitripennis* and *N. longicornis* females. Compounds in bold represent compounds that are part of peaks responsible for 50% dissimilarity in females.

| S.No. | Compound name | Average dissimilarity | Contribution % | Cumulative % | Mean Hosts with wasps (NV) | Mean Hosts with wasps (NL) |
|---|---|---|---|---|---|---|
| 1 | MeC33(5-) | 5.74 | 14.16 | 14.16 | 0.147 | 11.6 |
| 2 | **MeC35(15-;13-;11-)** | 4.06 | 10.02 | 24.18 | 5.95 | 14.1 |
| 3 | MeC35(7-) | 3.915 | 9.659 | 33.84 | 8.12 | 0.293 |
| 4 | C31 | 3.416 | 8.428 | 42.27 | 18.9 | 12.1 |
| 5 | MeC31(3-) | 2.676 | 6.603 | 48.87 | 3.79 | 9.14 |
| 6 | C33 | 2.268 | 5.596 | 54.47 | 4.54 | 0 |
| 7 | MeC35(17-) | 2.109 | 5.203 | 59.67 | 0 | 4.22 |
| 8 | DiMeC31(7,x-) | 1.831 | 4.518 | 64.19 | 3.9 | 0.242 |
| 9 | MeC31(15-;13-;11-) | 1.661 | 4.097 | 68.28 | 5.75 | 2.46 |
| 10 | MeC33(15-;13-;11-) | 1.496 | 3.691 | 71.97 | 12.7 | 15.7 |
| 11 | MeC29(3-) | 1.248 | 3.08 | 75.05 | 0 | 2.5 |
| 12 | DiMeC29(5-,x-) | 1.176 | 2.9 | 77.96 | 2.45 | 0.0957 |
| 13 | C29 | 0.7841 | 1.934 | 79.89 | 4.2 | 2.63 |
| 14 | DiMeC33(7,19-;7,23-) | 0.7327 | 1.808 | 81.7 | 0.48 | 1.95 |
| 15 | MeC31(7-) | 0.7088 | 1.749 | 83.45 | 7.08 | 5.67 |
| 16 | C33-ene_2 | 0.6285 | 1.551 | 85 | 0.102 | 1.36 |
| 17 | MeC31(5-) | 0.6248 | 1.542 | 86.54 | 3.36 | 2.11 |
| 18 | MeC37(15-;13-) | 0.5527 | 1.364 | 87.9 | 1.11 | 0 |
| 19 | C35 | 0.4801 | 1.185 | 89.09 | 1.37 | 2.33 |
| 20 | MeC33(3-) | 0.4618 | 1.139 | 90.23 | 0.924 | 0 |
| 21 | MeC35(5-;3-) | 0.4142 | 1.022 | 91.25 | 1.26 | 1.98 |
| 22 | DiMeC33(11,15-;11,21-) | 0.3462 | 0.8543 | 92.1 | 1.34 | 0.645 |
| 23 | MeC29(7-) | 0.3102 | 0.7654 | 92.87 | 0.943 | 0.323 |
| 24 | C30 | 0.3035 | 0.7488 | 93.62 | 0.933 | 0.326 |
| 25 | MeC33(7-) | 0.2929 | 0.7227 | 94.34 | 0.787 | 1.12 |
| 26 | MeC29(15-;13-;11-) | 0.2552 | 0.6295 | 94.97 | 1.3 | 0.795 |
| 27 | MeC29(5-) | 0.2111 | 0.5209 | 95.49 | 0.89 | 0.468 |
| 28 | MeC32(6-) | 0.2015 | 0.4972 | 95.99 | 0.636 | 0.233 |
| 29 | C33-ene_1 | 0.1596 | 0.3939 | 96.38 | 0.0791 | 0.398 |
| 30 | C32 | 0.1584 | 0.3908 | 96.77 | 0.746 | 0.429 |
| 31 | DiMeC33(3,17-;3,15-) | 0.1531 | 0.3778 | 97.15 | 1.5 | 1.29 |
| 32 | MeC27(15-;13-;11-;9-) | 0.1202 | 0.2965 | 97.45 | 0.634 | 0.394 |
| 33 | C26 | 0.1161 | 0.2863 | 97.73 | 0.232 | 0 |
| 34 | TetraMeC31(3,7,11,15-) | 0.1097 | 0.2706 | 98 | 0.486 | 0.267 |
| 35 | C34 | 0.1071 | 0.2641 | 98.27 | 0.455 | 0.669 |
| 36 | MeC30(7-;8-) | 0.1032 | 0.2546 | 98.52 | 0.206 | 0 |
| 37 | C27 | 0.1028 | 0.2537 | 98.78 | 0.755 | 0.58 |
| 38 | TetraMeC33(3,7,11,15-) | 0.1017 | 0.2508 | 99.03 | 0.124 | 0.327 |
| 39 | C28 | 0.09796 | 0.2417 | 99.27 | 0.196 | 0 |

Table S13 continued…

| | | | | | | |
|---|---|---|---|---|---|---|
| 40 | C25 | 0.07674 | 0.1893 | 99.46 | 0.361 | 0.207 |
| 41 | DiMeC31(7,11-) | 0.06619 | 0.1633 | 99.62 | 0.391 | 0.511 |
| 42 | MeC27(3-) | 0.05905 | 0.1457 | 99.77 | 0.468 | 0.375 |
| 43 | C31-ene-1 | 0.04265 | 0.1052 | 99.87 | 0.0853 | 0 |
| 44 | DiMeC29(3,11-;3,13-;3,15-) | 0.03238 | 0.07988 | 99.95 | 0.171 | 0.117 |
| 45 | C31-ene-2 | 0.01976 | 0.04875 | 100 | 0.139 | 0.102 |
| 46 | MeC25(3-) | 0 | 0 | 100 | 0 | 0 |

**Table S14:** Results of SIMPER analysis of the cuticular hydrocarbon profiles of hosts parasitised by *N. longicornis* and *N. oneida* females. Compounds in bold represent compounds that are part of peaks responsible for 50% dissimilarity in females.

| S.No. | Compound name | Average dissimilarity | Contribution % | Cumulative % | Mean Hosts with wasps (NL) | Mean Hosts with wasps (NO) |
|---|---|---|---|---|---|---|
| 1 | **MeC35(15-;13-;11-)** | 3.575 | 12.78 | 12.78 | 6.92 | 14.1 |
| 2 | **MeC33(3-)** | 2.799 | 10.01 | 22.79 | 5.6 | 0 |
| 3 | **MeC33(5-)** | 2.339 | 8.363 | 31.15 | 6.95 | 11.6 |
| 4 | MeC31(3-) | 1.549 | 5.539 | 36.69 | 6.05 | 9.14 |
| 5 | **C31** | 1.506 | 5.385 | 42.07 | 15.1 | 12.1 |
| 6 | **MeC31(7-)** | 1.329 | 4.752 | 46.83 | 3.01 | 5.67 |
| 7 | C35 | 1.165 | 4.164 | 50.99 | 0 | 2.33 |
| 8 | MeC33(15-;13-;11-) | 1.13 | 4.042 | 55.03 | 17.9 | 15.7 |
| 9 | DiMeC33(7,19-;7,23-) | 0.973 | 3.479 | 58.51 | 0 | 1.95 |
| 10 | MeC31(15-;13-;11-) | 0.918 | 3.282 | 61.79 | 3.21 | 2.46 |
| 11 | C33 | 0.8369 | 2.992 | 64.78 | 1.67 | 0 |
| 12 | DiMeC31(7,x-) | 0.7944 | 2.84 | 67.62 | 1.83 | 0.242 |
| 13 | TetraMeC33(3,7,11,15-) | 0.7475 | 2.672 | 70.3 | 1.82 | 0.327 |
| 14 | C33-ene_2 | 0.6797 | 2.43 | 72.73 | 0 | 1.36 |
| 15 | MeC31(5-) | 0.6475 | 2.315 | 75.04 | 3.41 | 2.11 |
| 16 | MeC27(15-;13-;11-;9-) | 0.5879 | 2.102 | 77.14 | 1.57 | 0.394 |
| 17 | MeC29(3-) | 0.5705 | 2.04 | 79.18 | 3.64 | 2.5 |
| 18 | MeC27(3-) | 0.5521 | 1.974 | 81.16 | 1.48 | 0.375 |
| 19 | C27 | 0.5084 | 1.818 | 82.98 | 1.6 | 0.58 |
| 20 | MeC29(15-;13-;11-) | 0.461 | 1.648 | 84.62 | 1.72 | 0.795 |
| 21 | C26 | 0.3925 | 1.403 | 86.03 | 0.785 | 0 |
| 22 | DiMeC33(3,17-;3,15-) | 0.3377 | 1.208 | 87.23 | 1.96 | 1.29 |
| 23 | C34 | 0.3345 | 1.196 | 88.43 | 0 | 0.669 |
| 24 | MeC35(17-) | 0.3283 | 1.174 | 89.6 | 3.57 | 4.22 |
| 25 | MeC35(5-;3-) | 0.3179 | 1.137 | 90.74 | 1.49 | 1.98 |
| 26 | C29 | 0.3085 | 1.103 | 91.84 | 2.03 | 2.63 |
| 27 | C30 | 0.2438 | 0.8717 | 92.72 | 0.813 | 0.326 |
| 28 | MeC33(7-) | 0.2362 | 0.8445 | 93.56 | 1.43 | 1.12 |
| 29 | C33-ene_1 | 0.1992 | 0.7122 | 94.27 | 0 | 0.398 |
| 30 | C32 | 0.1912 | 0.6836 | 94.96 | 0.812 | 0.429 |
| 31 | C25 | 0.1838 | 0.657 | 95.61 | 0.575 | 0.207 |
| 32 | MeC25(3-) | 0.1735 | 0.6203 | 96.23 | 0.347 | 0 |
| 33 | DiMeC33(11,15-;11,21-) | 0.1543 | 0.5517 | 96.78 | 0.337 | 0.645 |
| 34 | MeC35(7-) | 0.1465 | 0.5237 | 97.31 | 0 | 0.293 |
| 35 | DiMeC29(5-,x-) | 0.1437 | 0.5139 | 97.82 | 0.383 | 0.0957 |
| 36 | TetraMeC31(3,7,11,15-) | 0.1335 | 0.4774 | 98.3 | 0 | 0.267 |
| 37 | DiMeC31(7,11-) | 0.1195 | 0.4274 | 98.73 | 0.75 | 0.511 |
| 38 | MeC32(6-) | 0.09545 | 0.3413 | 99.07 | 0.213 | 0.233 |
| 39 | MeC29(5-) | 0.08489 | 0.3035 | 99.37 | 0.576 | 0.468 |
| 40 | MeC29(7-) | 0.06638 | 0.2373 | 99.61 | 0.437 | 0.323 |

Table S14 continued…

| | | | | | | |
|---|---|---|---|---|---|---|
| 41 | DiMeC29(3,11-;3,13-;3,15-) | 0.05853 | 0.2093 | 99.82 | 0 | 0.117 |
| 42 | C31-ene-2 | 0.05081 | 0.1816 | 100 | 0 | 0.102 |
| 43 | MeC37(15-;13-) | 0 | 0 | 100 | 0 | 0 |
| 44 | C31-ene-1 | 0 | 0 | 100 | 0 | 0 |
| 45 | MeC30(7-;8-) | 0 | 0 | 100 | 0 | 0 |
| 46 | C28 | 0 | 0 | 100 | 0 | 0 |

**Table S15:** Results of SIMPER analysis of the cuticular hydrocarbon profiles of hosts parasitised by *N. longicornis* and *N. giraulti* females. Compounds in bold represent compounds that are part of peaks responsible for 50% dissimilarity in females.

| S.No. | Compound name | Average dissimilarity | Contribution % | Cumulative % | Mean Hosts with wasps (NL) | Mean Hosts with wasps (NG) |
|---|---|---|---|---|---|---|
| 1 | MeC35(15-;13-;11-) | 5.02 | 14.93 | 14.93 | 14.1 | 4.03 |
| 2 | MeC33(5-) | 4.44 | 13.2 | 28.13 | 11.6 | 2.75 |
| 3 | TetraMeC33(3,7,11,15-) | 3.017 | 8.972 | 37.1 | 0.327 | 6.36 |
| 4 | MeC35(17-) | 2.294 | 6.822 | 43.92 | 4.22 | 8.64 |
| 5 | **MeC33(3-)** | 2.094 | 6.226 | 50.15 | 0 | 4.19 |
| 6 | C33 | 1.391 | 4.135 | 54.29 | 0 | 2.78 |
| 7 | MeC31(7-) | 1.187 | 3.53 | 57.82 | 5.67 | 3.29 |
| 8 | C35 | 1.165 | 3.463 | 61.28 | 2.33 | 0 |
| 9 | MeC31(3-) | 1.098 | 3.264 | 64.54 | 9.14 | 6.95 |
| 10 | C31 | 1.095 | 3.257 | 67.8 | 12.1 | 14.3 |
| 11 | MeC31(15-;13-;11-) | 1.091 | 3.244 | 71.04 | 2.46 | 4.25 |
| 12 | DiMeC33(7,19-;7,23-) | 0.973 | 2.893 | 73.94 | 1.95 | 0 |
| 13 | MeC31(5-) | 0.8894 | 2.645 | 76.58 | 2.11 | 3.89 |
| 14 | MeC33(15-;13-;11-) | 0.8513 | 2.531 | 79.11 | 15.7 | 17.4 |
| 15 | C33-ene_2 | 0.6797 | 2.021 | 81.14 | 1.36 | 0 |
| 16 | MeC35(5-;3-) | 0.5954 | 1.771 | 82.91 | 1.98 | 0.985 |
| 17 | TetraMeC31(3,7,11,15-) | 0.4952 | 1.473 | 84.38 | 0.267 | 1.26 |
| 18 | DiMeC33(11,15-;11,21-) | 0.4923 | 1.464 | 85.84 | 0.645 | 1.41 |
| 19 | DiMeC33(3,17-;3,15-) | 0.461 | 1.371 | 87.21 | 1.29 | 2.21 |
| 20 | DiMeC31(7,x-) | 0.3963 | 1.178 | 88.39 | 0.242 | 1.03 |
| 21 | MeC29(15-;13-;11-) | 0.3472 | 1.032 | 89.42 | 0.795 | 1.49 |
| 22 | C32 | 0.3449 | 1.026 | 90.45 | 0.429 | 1.12 |
| 23 | C34 | 0.3345 | 0.9948 | 91.44 | 0.669 | 0 |
| 24 | MeC33(7-) | 0.3177 | 0.9448 | 92.39 | 1.12 | 0.638 |
| 25 | C29 | 0.2991 | 0.8893 | 93.28 | 2.63 | 2.1 |
| 26 | MeC29(3-) | 0.2171 | 0.6454 | 93.92 | 2.5 | 2.4 |
| 27 | DiMeC31(7,11-) | 0.2029 | 0.6033 | 94.53 | 0.511 | 0.917 |
| 28 | C33-ene_1 | 0.1992 | 0.5924 | 95.12 | 0.398 | 0 |
| 29 | MeC27(3-) | 0.1971 | 0.586 | 95.71 | 0.375 | 0.769 |
| 30 | C30 | 0.1928 | 0.5733 | 96.28 | 0.326 | 0.711 |
| 31 | MeC27(15-;13-;11-;9-) | 0.1912 | 0.5687 | 96.85 | 0.394 | 0.742 |
| 32 | MeC32(6-) | 0.1658 | 0.493 | 97.34 | 0.233 | 0.565 |
| 33 | C27 | 0.1469 | 0.4368 | 97.78 | 0.58 | 0.822 |
| 34 | MeC35(7-) | 0.1465 | 0.4356 | 98.21 | 0.293 | 0 |
| 35 | MeC29(5-) | 0.1324 | 0.3936 | 98.61 | 0.468 | 0.732 |
| 36 | C26 | 0.1294 | 0.3848 | 98.99 | 0 | 0.259 |
| 37 | C25 | 0.09535 | 0.2835 | 99.27 | 0.207 | 0.398 |
| 38 | DiMeC29(3,11-;3,13-;3,15-) | 0.05853 | 0.174 | 99.45 | 0.117 | 0 |
| 39 | MeC25(3-) | 0.0579 | 0.1722 | 99.62 | 0 | 0.116 |

Table S15 continued…

| | | | | | | |
|---|---|---|---|---|---|---|
| 40 | MeC29(7-) | 0.055 | 0.1636 | 99.78 | 0.323 | 0.42 |
| 41 | C31-ene-2 | 0.05081 | 0.1511 | 99.94 | 0.102 | 0 |
| 42 | DiMeC29(5-,x-) | 0.02162 | 0.06429 | 100 | 0.0957 | 0.114 |
| 43 | MeC37(15-;13-) | 0 | 0 | 100 | 0 | 0 |
| 44 | C31-ene-1 | 0 | 0 | 100 | 0 | 0 |
| 45 | MeC30(7-;8-) | 0 | 0 | 100 | 0 | 0 |
| 46 | C28 | 0 | 0 | 100 | 0 | 0 |

**Table S16:** Results of SIMPER analysis of the cuticular hydrocarbon profiles of hosts parasitised by *N. vitripennis* and *N. giraulti* females. Compounds in bold represent compounds that are part of peaks responsible for 50% dissimilarity in hosts.

| S.No. | Compound name | Average dissimilarity | Contribution % | Cumulative % | Mean NV Female | Mean NG Female |
|---|---|---|---|---|---|---|
| 1 | **MeC33(15-;13-;11-)** | 5.277 | 13.2 | 13.2 | 12 | 1.82 |
| 2 | C29 | 4.973 | 12.44 | 25.63 | 12.2 | 2.66 |
| 3 | **MeC33(3-)** | 3.454 | 8.637 | 34.27 | 1.48 | 8.13 |
| 4 | C35 | 3.441 | 8.606 | 42.88 | 1.66 | 8.3 |
| 5 | **C31** | 2.903 | 7.261 | 50.14 | 21.2 | 15.6 |
| 6 | MeC35(15-;13-;11-) | 2.889 | 7.224 | 57.36 | 2.64 | 8.21 |
| 7 | MeC31(7-) | 2.343 | 5.859 | 63.22 | 9.24 | 4.73 |
| 8 | DiMeC33(11,15-;11,21-) | 2.338 | 5.848 | 69.07 | 1.66 | 6.16 |
| 9 | MeC31(15-;13-;11-) | 1.416 | 3.541 | 72.61 | 6.26 | 3.53 |
| 10 | MeC35(17-) | 1.349 | 3.373 | 75.98 | 1.53 | 4.13 |
| 11 | MeC35(5-;3-) | 1.336 | 3.342 | 79.33 | 0.732 | 3.31 |
| 12 | MeC29(7-) | 1.233 | 3.085 | 82.41 | 2.66 | 0.28 |
| 13 | MeC31(3-) | 1.08 | 2.7 | 85.11 | 2.61 | 4.69 |
| 14 | MeC37(15-;13-) | 0.5303 | 1.326 | 86.44 | 0 | 1.02 |
| 15 | C32 | 0.5126 | 1.282 | 87.72 | 0.415 | 1.4 |
| 16 | MeC29(3-) | 0.4982 | 1.246 | 88.96 | 1.7 | 0.744 |
| 17 | MeC29(5-) | 0.4758 | 1.19 | 90.15 | 1.46 | 0.543 |
| 18 | MeC32(6-) | 0.396 | 0.9902 | 91.14 | 0.988 | 0.225 |
| 19 | MeC33(5-) | 0.371 | 0.9279 | 92.07 | 0.715 | 0 |
| 20 | C27 | 0.364 | 0.9102 | 92.98 | 1.16 | 0.854 |
| 21 | DiMeC33(3,17-;3,15-) | 0.2426 | 0.6067 | 93.59 | 3.15 | 3.52 |
| 22 | DiMeC31(7,x-) | 0.1942 | 0.4856 | 94.08 | 0.463 | 0.0884 |
| 23 | C33-ene_1 | 0.1899 | 0.4749 | 94.55 | 0.457 | 0.0988 |
| 24 | C30 | 0.1818 | 0.4546 | 95 | 1.2 | 0.848 |
| 25 | MeC31(5-) | 0.177 | 0.4425 | 95.45 | 5.51 | 5.46 |
| 26 | MeC29(15-;13-;11-) | 0.1668 | 0.4171 | 95.86 | 0.742 | 0.42 |
| 27 | TetraMeC33(3,7,11,15-) | 0.1552 | 0.3881 | 96.25 | 0.331 | 0.63 |
| 28 | DiMeC33(7,19-;7,23-) | 0.129 | 0.3226 | 96.58 | 0.251 | 0.497 |
| 29 | C31-ene-1 | 0.1214 | 0.3036 | 96.88 | 0.29 | 0.0558 |
| 30 | MeC27(5-) | 0.1127 | 0.2819 | 97.16 | 0.294 | 0.0772 |
| 31 | C33 | 0.1089 | 0.2723 | 97.43 | 1.05 | 1.19 |
| 32 | C25 | 0.1088 | 0.2722 | 97.71 | 0.271 | 0.265 |
| 33 | C33-ene_2 | 0.09844 | 0.2462 | 97.95 | 0.211 | 0.401 |
| 34 | MeC30(7-;8-) | 0.09724 | 0.2432 | 98.19 | 0.252 | 0.0647 |
| 35 | MeC27(3-) | 0.0861 | 0.2153 | 98.41 | 0.125 | 0.279 |
| 36 | MeC33(7-) | 0.0799 | 0.1998 | 98.61 | 0.813 | 0.659 |
| 37 | MeC35(7-) | 0.07858 | 0.1965 | 98.81 | 0.501 | 0.354 |
| 38 | C34 | 0.07394 | 0.1849 | 98.99 | 0.353 | 0.496 |
| 39 | MeC25(5-) | 0.07006 | 0.1752 | 99.17 | 0.201 | 0.0662 |
| 40 | MeC25(3-) | 0.06324 | 0.1581 | 99.32 | 0.0259 | 0.139 |

Table S16 continued…

| | | | | | | |
|---|---|---|---|---|---|---|
| 41 | TetraMeC31(3,7,11,15-) | 0.05549 | 0.1388 | 99.46 | 0.164 | 0.0573 |
| 42 | DiMeC31(7,11-) | 0.04895 | 0.1224 | 99.59 | 0.408 | 0.319 |
| 43 | C31-ene-2 | 0.04012 | 0.1003 | 99.69 | 0.113 | 0.0355 |
| 44 | C28 | 0.03699 | 0.0925 | 99.78 | 0.128 | 0.0723 |
| 45 | DiMeC29(5-,x-) | 0.0277 | 0.06928 | 99.85 | 0.155 | 0.101 |
| 46 | MeC27(15-;13-;11-;9-) | 0.02273 | 0.05685 | 99.9 | 0.0224 | 0.057 |
| 47 | DiMeC29(3,11-;3,13-;3,15-) | 0.01836 | 0.04592 | 99.95 | 0.118 | 0.105 |
| 48 | MeC27(7-) | 0.01484 | 0.03712 | 99.99 | 0.0435 | 0.02 |
| 49 | C26 | 0.004965 | 0.01242 | 100 | 0.0147 | 0.0178 |
| 50 | MeC28(4-) | 0 | 0 | 100 | 0 | 0 |

**Table S17:** Results of SIMPER analysis of the cuticular hydrocarbon profiles of hosts parasitised by *N. vitripennis* and *N. oneida* females. Compounds in bold represent compounds that are part of peaks responsible for 50% dissimilarity in hosts.

| S.No. | Compound name | Average dissimilarity | Contribution % | Cumulative % | Mean NV Female | Mean NO Female |
|---|---|---|---|---|---|---|
| 1 | C29 | 4.623 | 13.43 | 13.43 | 12.2 | 3 |
| 2 | **C31** | 4.318 | 12.54 | 25.97 | 21.2 | 29.8 |
| 3 | **MeC33(3-)** | 3.476 | 10.1 | 36.07 | 1.48 | 8.43 |
| 4 | **MeC31(7-)** | 2.523 | 7.329 | 43.4 | 9.24 | 4.2 |
| 5 | **MeC33(5-)** | 2.495 | 7.247 | 50.65 | 0.715 | 5.7 |
| 6 | DiMeC31(7,11-) | 2.298 | 6.674 | 57.32 | 0.408 | 5 |
| 7 | DiMeC31(7,x-) | 1.488 | 4.322 | 61.64 | 0.463 | 3.44 |
| 8 | MeC29(7-) | 1.235 | 3.589 | 65.23 | 2.66 | 0.187 |
| 9 | MeC31(15-;13-;11-) | 1.203 | 3.495 | 68.73 | 6.26 | 3.85 |
| 10 | MeC33(15-;13-;11-) | 0.9764 | 2.836 | 71.56 | 12 | 13.9 |
| 11 | DiMeC33(3,17-;3,15-) | 0.9001 | 2.615 | 74.18 | 3.15 | 1.35 |
| 12 | MeC31(3-) | 0.8789 | 2.553 | 76.73 | 2.61 | 1.47 |
| 13 | C32 | 0.8397 | 2.439 | 79.17 | 0.415 | 2.02 |
| 14 | DiMeC33(11,15-;11,21-) | 0.8308 | 2.413 | 81.58 | 1.66 | 0 |
| 15 | MeC29(3-) | 0.7721 | 2.243 | 83.82 | 1.7 | 0.155 |
| 16 | MeC29(5-) | 0.611 | 1.775 | 85.6 | 1.46 | 0.237 |
| 17 | C27 | 0.5781 | 1.679 | 87.28 | 1.16 | 0 |
| 18 | MeC35(15-;13-;11-) | 0.5744 | 1.668 | 88.95 | 2.64 | 3.36 |
| 19 | MeC33(7-) | 0.4512 | 1.311 | 90.26 | 0.813 | 1.44 |
| 20 | MeC29(15-;13-;11-) | 0.3044 | 0.8841 | 91.14 | 0.742 | 0.133 |
| 21 | MeC35(17-) | 0.2671 | 0.7759 | 91.92 | 1.53 | 0.996 |
| 22 | MeC35(7-) | 0.2507 | 0.7281 | 92.65 | 0.501 | 0 |
| 23 | C33-ene_1 | 0.2287 | 0.6643 | 93.31 | 0.457 | 0 |
| 24 | C35 | 0.2 | 0.5811 | 93.89 | 1.66 | 1.41 |
| 25 | MeC32(6-) | 0.1727 | 0.5017 | 94.39 | 0.988 | 0.642 |
| 26 | C30 | 0.1622 | 0.4712 | 94.86 | 1.2 | 1.23 |
| 27 | MeC27(5-) | 0.1472 | 0.4274 | 95.29 | 0.294 | 0 |
| 28 | C31-ene-1 | 0.1449 | 0.4209 | 95.71 | 0.29 | 0 |
| 29 | MeC31(5-) | 0.14 | 0.4068 | 96.12 | 5.51 | 5.79 |
| 30 | C33 | 0.1369 | 0.3976 | 96.52 | 1.05 | 0.78 |
| 31 | C25 | 0.1354 | 0.3933 | 96.91 | 0.271 | 0 |
| 32 | MeC35(5-;3-) | 0.1263 | 0.3669 | 97.28 | 0.732 | 0.61 |
| 33 | MeC30(7-;8-) | 0.1261 | 0.3662 | 97.64 | 0.252 | 0 |
| 34 | C33-ene_2 | 0.1054 | 0.3061 | 97.95 | 0.211 | 0 |
| 35 | MeC25(5-) | 0.1005 | 0.292 | 98.24 | 0.201 | 0 |
| 36 | TetraMeC31(3,7,11,15-) | 0.0821 | 0.2385 | 98.48 | 0.164 | 0 |
| 37 | TetraMeC33(3,7,11,15-) | 0.0738 | 0.2144 | 98.69 | 0.331 | 0.184 |
| 38 | C34 | 0.06986 | 0.2029 | 98.9 | 0.353 | 0.214 |
| 39 | C28 | 0.06379 | 0.1853 | 99.08 | 0.128 | 0 |
| 40 | MeC27(3-) | 0.06271 | 0.1822 | 99.27 | 0.125 | 0 |

Table S17 continued…

| | | | | | | |
|---|---|---|---|---|---|---|
| 41 | DiMeC29(3,11-;3,13-;3,15-) | 0.0592 | 0.172 | 99.44 | 0.118 | 0 |
| 42 | C31-ene-2 | 0.0564 | 0.1638 | 99.6 | 0.113 | 0 |
| 43 | DiMeC29(5-,x-) | 0.04308 | 0.1251 | 99.73 | 0.155 | 0.0684 |
| 44 | DiMeC33(7,19-;7,23-) | 0.04102 | 0.1192 | 99.85 | 0.251 | 0.315 |
| 45 | MeC27(7-) | 0.02176 | 0.06321 | 99.91 | 0.0435 | 0 |
| 46 | MeC25(3-) | 0.01297 | 0.03767 | 99.95 | 0.0259 | 0 |
| 47 | MeC27(15-;13-;11-;9-) | 0.01123 | 0.03261 | 99.98 | 0.0224 | 0 |
| 48 | C26 | 0.00733 | 0.02129 | 100 | 0.0147 | 0 |
| 49 | MeC37(15-;13-) | 0 | 0 | 100 | 0 | 0 |
| 50 | MeC28(4-) | 0 | 0 | 100 | 0 | 0 |

**Table S18:** Results of SIMPER analysis of the cuticular hydrocarbon profiles of hosts parasitised by *N. vitripennis* and *N. longicornis* females. Compounds in bold represent compounds that are part of peaks responsible for 50% dissimilarity in hosts.

| S.No. | Compound name | Average dissimilarity | Contribution % | Cumulative % | Mean NV Female | Mean NL Female |
|---|---|---|---|---|---|---|
| 1 | **MeC35(15-;13-;11-)** | 2.215 | 11.15 | 11.15 | 2.64 | 7.07 |
| 2 | TetraMeC31(3,7,11,15-) | 1.756 | 8.838 | 19.99 | 0.164 | 3.68 |
| 3 | MeC31(15-;13-;11-) | 1.678 | 8.445 | 28.43 | 6.26 | 2.9 |
| 4 | MeC33(15-;13-;11-) | 1.413 | 7.11 | 35.54 | 12 | 14.8 |
| 5 | DiMeC33(3,17-;3,15-) | 1.107 | 5.569 | 41.11 | 3.15 | 0.932 |
| 6 | MeC31(5-) | 1.05 | 5.285 | 46.39 | 5.51 | 3.41 |
| 7 | DiMeC31(7,11-) | 0.8066 | 4.059 | 50.45 | 0.408 | 2.02 |
| 8 | MeC29(7-) | 0.7105 | 3.575 | 54.03 | 2.66 | 1.24 |
| 9 | DiMeC33(11,15-;11,21-) | 0.6423 | 3.232 | 57.26 | 1.66 | 0.377 |
| 10 | MeC33(3-) | 0.6116 | 3.078 | 60.34 | 1.48 | 2.7 |
| 11 | C34 | 0.6037 | 3.038 | 63.38 | 0.353 | 1.56 |
| 12 | C31 | 0.5569 | 2.802 | 66.18 | 21.2 | 21.2 |
| 13 | MeC29(5-) | 0.5284 | 2.659 | 68.84 | 1.46 | 0.403 |
| 14 | C33 | 0.5267 | 2.651 | 71.49 | 1.05 | 0 |
| 15 | C29 | 0.4695 | 2.362 | 73.85 | 12.2 | 11.3 |
| 16 | MeC32(6-) | 0.4343 | 2.186 | 76.04 | 0.988 | 0.119 |
| 17 | MeC29(3-) | 0.4099 | 2.063 | 78.1 | 1.7 | 2.52 |
| 18 | C32 | 0.4059 | 2.042 | 80.14 | 0.415 | 1.23 |
| 19 | MeC31(3-) | 0.4022 | 2.024 | 82.17 | 2.61 | 1.81 |
| 20 | C27 | 0.3137 | 1.578 | 83.74 | 1.16 | 0.529 |
| 21 | DiMeC33(7,19-;7,23-) | 0.2868 | 1.443 | 85.19 | 0.251 | 0.731 |
| 22 | MeC31(7-) | 0.2528 | 1.272 | 86.46 | 9.24 | 9.05 |
| 23 | MeC35(7-) | 0.2507 | 1.261 | 87.72 | 0.501 | 0 |
| 24 | MeC35(17-) | 0.2389 | 1.202 | 88.92 | 1.53 | 2.01 |
| 25 | MeC33(7-) | 0.2174 | 1.094 | 90.02 | 0.813 | 0.378 |
| 26 | C33-ene_1 | 0.2084 | 1.049 | 91.07 | 0.457 | 0.241 |
| 27 | C30 | 0.1895 | 0.9538 | 92.02 | 1.2 | 1.58 |
| 28 | MeC30(7-;8-) | 0.1877 | 0.9445 | 92.96 | 0.252 | 0.627 |
| 29 | TetraMeC33(3,7,11,15-) | 0.1657 | 0.8337 | 93.8 | 0.331 | 0 |
| 30 | MeC33(5-) | 0.1465 | 0.7371 | 94.54 | 0.715 | 0.422 |
| 31 | DiMeC31(7,x-) | 0.146 | 0.7347 | 95.27 | 0.463 | 0.171 |
| 32 | C25 | 0.1347 | 0.6781 | 95.95 | 0.271 | 0.273 |
| 33 | MeC29(15-;13-;11-) | 0.1096 | 0.5516 | 96.5 | 0.742 | 0.961 |
| 34 | C35 | 0.1073 | 0.5399 | 97.04 | 1.66 | 1.54 |
| 35 | MeC35(5-;3-) | 0.09006 | 0.4532 | 97.49 | 0.732 | 0.912 |
| 36 | MeC25(5-) | 0.06839 | 0.3441 | 97.84 | 0.201 | 0.0643 |
| 37 | C31-ene-1 | 0.06206 | 0.3123 | 98.15 | 0.29 | 0.207 |
| 38 | DiMeC29(3,11-;3,13-;3,15-) | 0.0592 | 0.2979 | 98.45 | 0.118 | 0 |
| 39 | C26 | 0.05627 | 0.2831 | 98.73 | 0.0147 | 0.127 |
| 40 | C33-ene_2 | 0.05625 | 0.2831 | 99.01 | 0.211 | 0.0982 |
| 41 | C31-ene-2 | 0.04681 | 0.2355 | 99.25 | 0.113 | 0.0192 |

Table S18 continued…

| | | | | | | |
|---|---|---|---|---|---|---|
| 42 | MeC27(3-) | 0.03425 | 0.1723 | 99.42 | 0.125 | 0.0755 |
| 43 | MeC27(5-) | 0.02846 | 0.1432 | 99.56 | 0.294 | 0.299 |
| 44 | MeC25(3-) | 0.01852 | 0.09319 | 99.66 | 0.0259 | 0.063 |
| 45 | DiMeC29(5-,x-) | 0.01678 | 0.08442 | 99.74 | 0.155 | 0.121 |
| 46 | C28 | 0.01342 | 0.06751 | 99.81 | 0.128 | 0.147 |
| 47 | MeC27(15-;13-;11-;9-) | 0.01289 | 0.06484 | 99.87 | 0.0224 | 0.0482 |
| 48 | MeC27(7-) | 0.01263 | 0.06358 | 99.94 | 0.0435 | 0.0206 |
| 49 | MeC28(4-) | 0.01239 | 0.06233 | 100 | 0 | 0.0248 |
| 50 | MeC37(15-;13-) | 0 | 0 | 100 | 0 | 0 |

**Table S19:** Results of SIMPER analysis of the cuticular hydrocarbon profiles of hosts parasitised by *N. longicornis* and *N. oneida* females. Compounds in bold represent compounds that are part of peaks responsible for 50% dissimilarity in hosts.

| S.No. | Compound name | Average dissimilarity | Contribution % | Cumulative % | Mean NL Female | Mean NO Female |
|---|---|---|---|---|---|---|
| 1 | **C31** | 4.329 | 12.59 | 12.59 | 21.2 | 29.8 |
| 2 | C29 | 4.153 | 12.08 | 24.67 | 11.3 | 3 |
| 3 | **MeC33(3-)** | 2.864 | 8.329 | 33 | 2.7 | 8.43 |
| 4 | **MeC33(5-)** | 2.641 | 7.681 | 40.68 | 0.422 | 5.7 |
| 5 | **MeC31(7-)** | 2.426 | 7.054 | 47.73 | 9.05 | 4.2 |
| 6 | **MeC35(15-;13-;11-)** | 1.854 | 5.392 | 53.12 | 7.07 | 3.36 |
| 7 | TetraMeC31(3,7,11,15-) | 1.838 | 5.346 | 58.47 | 3.68 | 0 |
| 8 | DiMeC31(7,x-) | 1.634 | 4.751 | 63.22 | 0.171 | 3.44 |
| 9 | DiMeC31(7,11-) | 1.491 | 4.336 | 67.55 | 2.02 | 5 |
| 10 | MeC31(5-) | 1.187 | 3.453 | 71.01 | 3.41 | 5.79 |
| 11 | MeC29(3-) | 1.182 | 3.437 | 74.44 | 2.52 | 0.155 |
| 12 | MeC31(3-) | 0.7448 | 2.166 | 76.61 | 1.81 | 1.47 |
| 13 | C32 | 0.7044 | 2.048 | 78.66 | 1.23 | 2.02 |
| 14 | C34 | 0.6735 | 1.958 | 80.62 | 1.56 | 0.214 |
| 15 | MeC33(15-;13-;11-) | 0.6364 | 1.851 | 82.47 | 14.8 | 13.9 |
| 16 | MeC33(7-) | 0.5375 | 1.563 | 84.03 | 0.378 | 1.44 |
| 17 | MeC29(7-) | 0.5249 | 1.526 | 85.56 | 1.24 | 0.187 |
| 18 | MeC35(17-) | 0.506 | 1.471 | 87.03 | 2.01 | 0.996 |
| 19 | MeC31(15-;13-;11-) | 0.4751 | 1.382 | 88.41 | 2.9 | 3.85 |
| 20 | MeC29(15-;13-;11-) | 0.414 | 1.204 | 89.61 | 0.961 | 0.133 |
| 21 | C33 | 0.3899 | 1.134 | 90.75 | 0 | 0.78 |
| 22 | MeC30(7-;8-) | 0.3137 | 0.9123 | 91.66 | 0.627 | 0 |
| 23 | DiMeC33(7,19-;7,23-) | 0.2762 | 0.8032 | 92.46 | 0.731 | 0.315 |
| 24 | C27 | 0.2645 | 0.7691 | 93.23 | 0.529 | 0 |
| 25 | MeC32(6-) | 0.2616 | 0.7607 | 93.99 | 0.119 | 0.642 |
| 26 | C30 | 0.214 | 0.6224 | 94.61 | 1.58 | 1.23 |
| 27 | DiMeC33(3,17-;3,15-) | 0.2067 | 0.601 | 95.22 | 0.932 | 1.35 |
| 28 | DiMeC33(11,15-;11,21-) | 0.1885 | 0.5482 | 95.76 | 0.377 | 0 |
| 29 | MeC29(5-) | 0.1687 | 0.4905 | 96.25 | 0.403 | 0.237 |
| 30 | MeC35(5-;3-) | 0.1631 | 0.4742 | 96.73 | 0.912 | 0.61 |
| 31 | C35 | 0.1533 | 0.4457 | 97.17 | 1.54 | 1.41 |
| 32 | MeC27(5-) | 0.1493 | 0.434 | 97.61 | 0.299 | 0 |
| 33 | C25 | 0.1364 | 0.3967 | 98 | 0.273 | 0 |
| 34 | C33-ene_1 | 0.1203 | 0.3499 | 98.35 | 0.241 | 0 |
| 35 | C31-ene-1 | 0.1035 | 0.3009 | 98.66 | 0.207 | 0 |
| 36 | TetraMeC33(3,7,11,15-) | 0.09187 | 0.2671 | 98.92 | 0 | 0.184 |
| 37 | C28 | 0.07367 | 0.2142 | 99.14 | 0.147 | 0 |
| 38 | C26 | 0.0636 | 0.1849 | 99.32 | 0.127 | 0 |
| 39 | C33-ene_2 | 0.04912 | 0.1428 | 99.46 | 0.0982 | 0 |

Table S19 continued…

| | | | | | | |
|---|---|---|---|---|---|---|
| 40 | MeC27(3-) | 0.03774 | 0.1098 | 99.57 | 0.0755 | 0 |
| 41 | MeC25(5-) | 0.03215 | 0.09348 | 99.67 | 0.0643 | 0 |
| 42 | MeC25(3-) | 0.03149 | 0.09155 | 99.76 | 0.063 | 0 |
| 43 | DiMeC29(5-,x-) | 0.0263 | 0.07649 | 99.84 | 0.121 | 0.0684 |
| 44 | MeC27(15-;13-;11-;9-) | 0.02411 | 0.07011 | 99.91 | 0.0482 | 0 |
| 45 | MeC28(4-) | 0.01239 | 0.03602 | 99.94 | 0.0248 | 0 |
| 46 | MeC27(7-) | 0.01029 | 0.02993 | 99.97 | 0.0206 | 0 |
| 47 | C31-ene-2 | 0.009588 | 0.02788 | 100 | 0.0192 | 0 |
| 48 | MeC37(15-;13-) | 0 | 0 | 100 | 0 | 0 |
| 49 | MeC35(7-) | 0 | 0 | 100 | 0 | 0 |
| 50 | DiMeC29(3,11-;3,13-;3,15-) | 0 | 0 | 100 | 0 | 0 |

**Table S20:** Results of SIMPER analysis of the cuticular hydrocarbon profiles of hosts parasitised by *N. longicornis* and *N. giraulti* females. Compounds in bold represent compounds that are part of peaks responsible for 50% dissimilarity in hosts.

| S.No. | Compound name | Average dissimilarity | Contribution % | Cumulative % | Mean NL Female | Mean NG Female |
|---|---|---|---|---|---|---|
| 1 | MeC33(15-;13-;11-) | 6.743 | 15.78 | 15.78 | 14.8 | 1.82 |
| 2 | C29 | 4.486 | 10.5 | 26.29 | 11.3 | 2.66 |
| 3 | C35 | 3.503 | 8.2 | 34.49 | 1.54 | 8.3 |
| 4 | DiMeC33(11,15-;11,21-) | 3.005 | 7.034 | 41.52 | 0.377 | 6.16 |
| 5 | C31 | 2.892 | 6.769 | 48.29 | 21.2 | 15.6 |
| 6 | **MeC33(3-)** | 2.819 | 6.599 | 54.89 | 2.7 | 8.13 |
| 7 | MeC31(7-) | 2.242 | 5.247 | 60.13 | 9.05 | 4.73 |
| 8 | TetraMeC31(3,7,11,15-) | 1.878 | 4.396 | 64.53 | 3.68 | 0.0573 |
| 9 | MeC31(3-) | 1.497 | 3.504 | 68.03 | 1.81 | 4.69 |
| 10 | DiMeC33(3,17-;3,15-) | 1.341 | 3.139 | 71.17 | 0.932 | 3.52 |
| 11 | MeC35(5-;3-) | 1.243 | 2.91 | 74.08 | 0.912 | 3.31 |
| 12 | MeC35(17-) | 1.101 | 2.577 | 76.66 | 2.01 | 4.13 |
| 13 | MeC31(5-) | 1.06 | 2.482 | 79.14 | 3.41 | 5.46 |
| 14 | MeC29(3-) | 0.9214 | 2.157 | 81.3 | 2.52 | 0.744 |
| 15 | DiMeC31(7,11-) | 0.8831 | 2.067 | 83.37 | 2.02 | 0.319 |
| 16 | MeC35(15-;13-;11-) | 0.6735 | 1.576 | 84.94 | 7.07 | 8.21 |
| 17 | C33 | 0.6176 | 1.446 | 86.39 | 0 | 1.19 |
| 18 | C34 | 0.5525 | 1.293 | 87.68 | 1.56 | 0.496 |
| 19 | MeC37(15-;13-) | 0.5303 | 1.241 | 88.92 | 0 | 1.02 |
| 20 | MeC29(7-) | 0.4961 | 1.161 | 90.08 | 1.24 | 0.28 |
| 21 | MeC31(15-;13-;11-) | 0.4545 | 1.064 | 91.15 | 2.9 | 3.53 |
| 22 | C30 | 0.3785 | 0.8859 | 92.03 | 1.58 | 0.848 |
| 23 | TetraMeC33(3,7,11,15-) | 0.3271 | 0.7657 | 92.8 | 0 | 0.63 |
| 24 | C27 | 0.3008 | 0.704 | 93.5 | 0.529 | 0.854 |
| 25 | MeC30(7-;8-) | 0.292 | 0.6835 | 94.19 | 0.627 | 0.0647 |
| 26 | MeC29(15-;13-;11-) | 0.2805 | 0.6566 | 94.84 | 0.961 | 0.42 |
| 27 | DiMeC33(7,19-;7,23-) | 0.2551 | 0.5973 | 95.44 | 0.731 | 0.497 |
| 28 | MeC29(5-) | 0.2278 | 0.5332 | 95.97 | 0.403 | 0.543 |
| 29 | MeC33(5-) | 0.219 | 0.5127 | 96.49 | 0.422 | 0 |
| 30 | MeC35(7-) | 0.1836 | 0.4297 | 96.92 | 0 | 0.354 |
| 31 | C33-ene_2 | 0.1568 | 0.3671 | 97.28 | 0.0982 | 0.401 |
| 32 | C25 | 0.1486 | 0.3479 | 97.63 | 0.273 | 0.265 |
| 33 | MeC33(7-) | 0.1458 | 0.3413 | 97.97 | 0.378 | 0.659 |
| 34 | MeC27(5-) | 0.1149 | 0.269 | 98.24 | 0.299 | 0.0772 |
| 35 | MeC27(3-) | 0.1052 | 0.2463 | 98.49 | 0.0755 | 0.279 |
| 36 | C33-ene_1 | 0.09269 | 0.217 | 98.71 | 0.241 | 0.0988 |
| 37 | C32 | 0.09138 | 0.2139 | 98.92 | 1.23 | 1.4 |
| 38 | C31-ene-1 | 0.07842 | 0.1836 | 99.1 | 0.207 | 0.0558 |
| 39 | MeC25(3-) | 0.06071 | 0.1421 | 99.25 | 0.063 | 0.139 |
| 40 | C26 | 0.05679 | 0.1329 | 99.38 | 0.127 | 0.0178 |

Table S20 continued…

| | | | | | | |
|---|---|---|---|---|---|---|
| 41 | MeC32(6-) | 0.05475 | 0.1282 | 99.51 | 0.119 | 0.225 |
| 42 | DiMeC29(3,11-;3,13-;3,15-) | 0.05422 | 0.1269 | 99.63 | 0 | 0.105 |
| 43 | DiMeC31(7,x-) | 0.04267 | 0.09989 | 99.73 | 0.171 | 0.0884 |
| 44 | C28 | 0.03975 | 0.09305 | 99.83 | 0.147 | 0.0723 |
| 45 | MeC27(15-;13-;11-;9-) | 0.01966 | 0.04603 | 99.87 | 0.0482 | 0.057 |
| 46 | MeC25(5-) | 0.01656 | 0.03876 | 99.91 | 0.0643 | 0.0662 |
| 47 | MeC28(4-) | 0.01285 | 0.03009 | 99.94 | 0.0248 | 0 |
| 48 | DiMeC29(5-,x-) | 0.0109 | 0.0255 | 99.97 | 0.121 | 0.101 |
| 49 | C31-ene-2 | 0.008458 | 0.0198 | 99.99 | 0.0192 | 0.0355 |
| 50 | MeC27(7-) | 0.005793 | 0.01356 | 100 | 0.0206 | 0.02 |

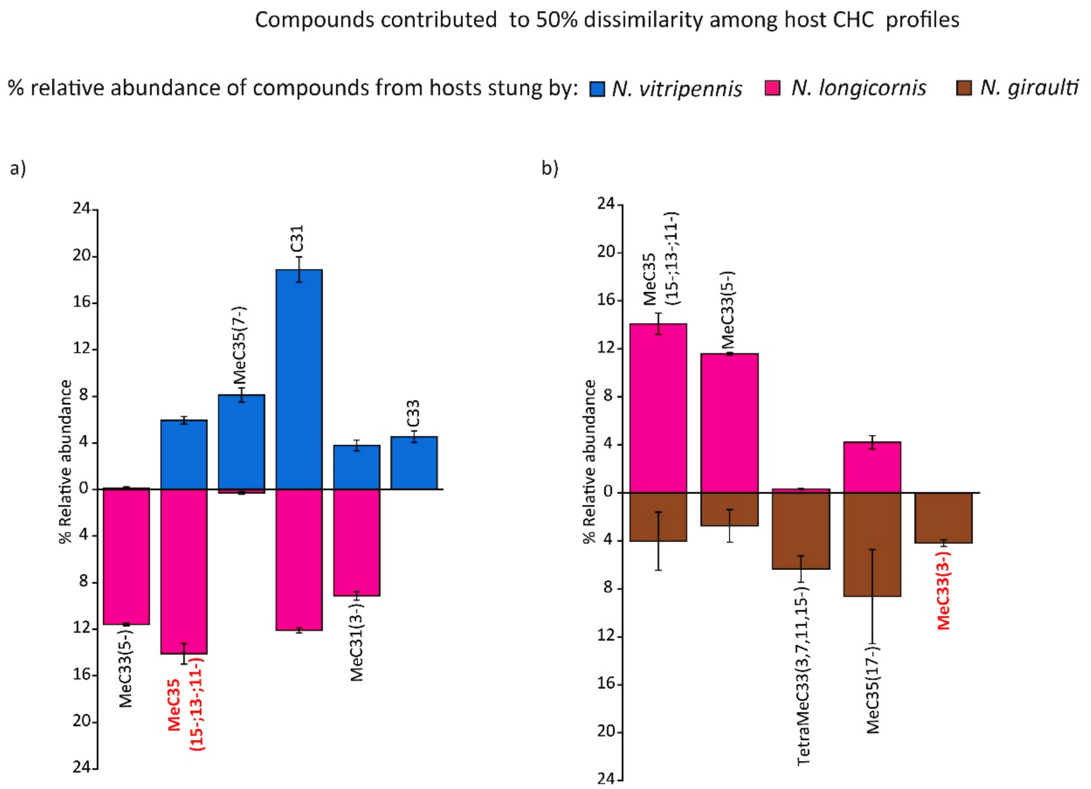


**Figure S1**: Relative abundance (%) of compounds that cumulatively contributed to 50% of dissimilarity between CHC profiles of parasitised hosts, which were distinguished by males: Comparison of compounds between a) hosts parasitised by *N. vitripennis* and *N. longicornis,* b) *N. longicornis* and *N. giraulti*. Compounds titled in red also contribute to dissimilarity in host profiles of the corresponding species.
